# *In vitro* teratogenicity testing using a 3D, embryo-like gastruloid system

**DOI:** 10.1101/2021.03.30.437698

**Authors:** Veronika Mantziou, Peter Baillie-Benson, Manuela Jaklin, Stefan Kustermann, Alfonso Martinez Arias, Naomi Moris

## Abstract

Pharmaceuticals that are intended for use in patients of childbearing potential need to be tested for teratogenicity before marketing. Several pharmaceutical companies use animal-free *in vitro* models which allow a more rapid selection of lead compounds and contribute to 3Rs principles (‘replace, reduce and refine’) by streamlining the selection of promising compounds that are submitted to further regulatory studies in animals. Currently available *in vitro* models typically rely on adherent monolayer cultures or disorganized 3D structures, both of which lack the spatiotemporal and morphological context of the developing embryo. A newly developed 3D ‘gastruloid’ model has the potential to achieve a more reliable prediction of teratogenicity by providing a robust recapitulation of gastrulation-like events alongside morphological coordination at relatively high-throughput. In this first proof-of-concept study, we used both mouse and human gastruloids to examine a panel of seven reference compounds, with associated *in vivo* data and known teratogenic risk, to quantitatively assess *in vitro* teratogenicity. We observed several gross morphological effects, including significantly reduced elongation or decreased size of the gastruloids, upon exposure to several of the reference compounds. We also observed aberrant gene expression using fluorescent reporters, including *SOX2, BRA*, and *SOX17*, suggestive of multi-lineage differentiation defects and disrupted axial patterning. Finally, we saw that gastruloids recapitulated some of the known *in vivo* species-specific susceptibilities between their mouse and human counterparts. We therefore suggest that gastruloids represent a powerful tool for teratogenicity assessment by enabling relevant physiological recapitulation of early embryonic development, demonstrating their use as a novel *in vitro* teratogenic model system.

## 1. Introduction

Developing embryos are highly susceptible to any harmful exposure to teratogenic substances, due to incomplete epithelial barrier functions and detoxification capacity, especially in the first trimester of pregnancy [1, 2]. Teratogens may cause congenital abnormalities to occur, leading to lifelong physical or functional impairment. However, the underlying mechanisms of many teratogens are still not well described and further investigations are necessary to enable us to understand, let alone predict, which disruptive effects might cause birth defects.

Particularly in the application of pharmaceuticals intended for women of childbearing potential, it is crucial to know which substances may bear teratogenic risks. Therefore, promising drug candidates must be screened for any developmental or fetal toxicity prior to clinical application. This involves testing in pregnant animals in order to predict the likely effect of compound exposure at different concentrations. However, a number of high-profile cases have suggested that model organisms like the mouse are often unable to accurately predict human teratogenicity, such as thalidomide [3, 4]. As a result, Developmental and Reproductive Toxicology (DART) studies now incorporate testing on several species, including non-rodents such as rabbits or nonhuman primates, in order to have increased confidence in the safety of a new compound [5].

An alternative option, fast gaining traction in pharmaceutical companies, is to utilize *in vitro* systems, such as pluripotent stem cells, to design assays for teratogenicity assessment [6]. The benefit of such systems is threefold: they facilitate human-specific predictions by using human cell lines; they reduce the number of animals required for research, thus contributing to a 3Rs principle (to ‘replace, reduce and refine’ animal use in research) and they allow for medium- to high-throughput analyses earlier in the drug development pipeline. As such, *in vitro* model systems that predict human teratogenicity represent a powerful new approach in pharmaceutical development.

Recently, several *in vitro* models have been developed and applied to the evaluation of developmental toxicants [7–10]. However, most of these models are established as adherent monolayer cultures, which are lacking an essential micro-physiological environment and the readouts of assays often only target a single germ layer. Therefore, commonly available *in vitro* systems do not fully reflect all the relevant features of spatial and temporal patterning or multilineage differentiation during gastrulation [11, 12]. Instead, a more biologically relevant model that better recreates the native processes of early embryonic development, could reveal new mechanistic insights into disruptive events during embryogenesis and might improve prediction of teratogenicity.

Recent work has described a pluripotent stem cell (PSC)-based system that utilizes the self-organizing potential of PSCs to generate three-dimensional structures that recapitulate elements of the early embryo [13–15]. These are known as ‘gastruloids’, as they mirror some of the events of gastrulation; where the emergence of the three germ layers is coordinated along anterior-posterior, dorsal-ventral and medio-lateral axes to generate the elements of the body plan, from which the various tissues and organs will emerge. Studies have shown that these stem cell aggregates progressively break symmetry, undergo axial elongation and differentiate to all three germ layers (mesoderm, ectoderm and endoderm) in a manner that is spatially and temporally similar to the embryo. However, these structures do not contain anterior neural (brain) or extraembryonic cell types, meaning they do not have full organismal potential [16]. Recently, this work with mouse PSCs has been extended to the generation of gastruloids from human Embryonic Stem Cells (ESCs) [15]. These so-called, ‘human gastruloids’, recapitulate many features of the developing mammalian embryo, and may bring unique insights into human development with experimental tractability that overcomes several of the ethical and technical limitations of research on human embryos.

Gastruloids have reproducible morphological changes, germ layer proportions and spatiotemporal organization of gene expression that are easy to visualize and quantify. These are key benefits over traditional adherent cell methods or embryoid body (EB) cultures. They therefore offer an experimentally tractable system to explore the effect of a range of perturbations that may affect cell lineage differentiation, signaling, morphology, cell viability and growth [13, 15].

Here, we examine whether mouse and human gastruloids can have further application as an effective teratogenicity assay. Using a small reference panel of compounds; including *all-trans* retinoic acid, valproic acid, bosentan, thalidomide, phenytoin, ibuprofen and penicillin G; we explore gastruloid sensitivity to different concentrations and exposures. These are then compared between mouse and human gastruloids, against existing data from gold-standard animal models and compared to known human teratological status. We use a range of outputs including morphological shape descriptors, marker gene expression and cytotoxicity to qualitatively and quantitatively assess the effect of chemical exposure on gastruloids. Distribution of germ layer representative reporter gene expression of *SOX2* (neuroectoderm), *SOX17* (endoderm) and *BRA* (mesoderm) within human gastruloids, and *T/Bra* in mouse gastruloids, enabled us to examine the correct generation of germ layer lineages and the extent of axial polarity. Furthermore, we compared the data with those from existing 3D mouse and human *in vitro* models to see whether the gastruloid systems are able to advance the physiological relevance compared to systems based on morphologically less structured embryoid bodies [7, 17, 18].

## 2. Materials and Methods

### 2.1. Mouse Embryonic Stem Cell Culture

Mouse embryonic stem cells were maintained as previously described, on gelatin-precoated tissue culture plastic in ES+LIF (leukemia inhibitory factor) medium [19]. Cells were passaged at a density of 8 x 10^3^ cells/cm^2^ every second day, with approximately two-thirds of the culture medium exchanged for fresh ES+LIF medium on the intervening days. The cell lines used in this study were E14Tg2A [20] and T/Bra::GFP [21]. They were maintained in culture for at least two passages post-thawing prior to experimental use and they were propagated for no more than 30 passages *in vitro*. All cell counting during routine maintenance and experimentation was performed with an ORFLO Moxi Z mini automated cell counter (ORFLO Technologies, MXZ002).

### 2.2. Mouse Gastruloid Culture

Gastruloids were prepared by aggregating 300 mouse embryonic stem cells in 40 μl droplets of N2B27 (NDiff^®^ 227, Takara Bio Inc. Y40002) per well of U-bottomed 96-well plates (Greiner 650185) as described previously [13]. The following modifications were made to the protocol. Cell cultures were plated into new flasks at a density of 8 x 10^3^ cells/cm^2^ in ES+LIF medium two days prior to preparing gastruloids. The culture medium was changed fully to 2i+LIF medium the following day as a 24-hour pre-treatment.

The compounds were administered to the gastruloids on plating and daily after aggregation was complete (at 0 h, 48 h, 72 h and 96 h post-plating). The compounds were prepared fresh from frozen stocks in N2B27 culture medium at each time point.

### 2.3. Human Embryonic Stem Cell Culture

The human cell line used in this study was the ES cell line RUES2-GLR (mCit–*SOX2*, mCerulean–*BRA*, tdTomato–*SOX17*) [22]. All cells were cultured in humidified incubators at 37°C and 5% CO_2_. Human ES cells were cultured routinely in Nutristem hPSC XF medium (Biological Industries, 05-100-1A) on 0.5 μg/cm^2^ Vitronectin-coated plates (Gibco, A14700). Cells were passaged using 0.5 mM EDTA in phosphate-buffered saline without Mg^2+^ or Ca^2+^ (PBS^-/-^) (Invitrogen, 15575-038).

### 2.4. Human Gastruloid Culture

Before human gastruloid culture, cells were passaged to single cell suspension and plated to achieve 65,000 cells per well of a 6-well plate in Nutristem supplemented with 1:2,000 Y-27632 (ROCK inhibitor; Sigma Aldrich, Y0503) [23]. Adherent cultures were pre-treated in Nutristem supplemented with 3.25 μM CHIR99021 (Chiron; Tocris Biosciences, 4423) and either 0.2% DMSO (vehicle), water (vehicle for penicillin G) or the different compound concentrations on the fourth day of adherent culture. After pre-treatment for 24 h, cells were dissociated using 0.5 mM EDTA in PBS^-/-^ (Invitrogen, 15575-038), washed in PBS^-/-^ and reaggregated in basal differentiation medium, Essential 6 (E6; Thermo Fisher Scientific, A15165-01), supplemented with 1:2,000 Y-27632 (ROCK inhibitor), 0.5 μM Chiron and either 0.2% DMSO, water (vehicles) or the different compound concentrations. Cell numbers were determined using an automated cell counter (Moxi Z Mini, ORFLO Technologies, MXZ002) and 400 cells per 40 μl were added to each well of an ultra-low-adherence 96-well plate (CellStar, 650970). The cell suspension was centrifuged using a benchtop plate centrifuge (Eppendorf) at 700 rpm at room temperature for 2 min. The following day, 150 μl fresh E6 medium was added to each well. The medium was exchanged for fresh E6 medium daily after this time point (150 μl per well).

### 2.5. Embryoid body based in vitro systems

To evaluate the data from the gastruloid models, described protocols of the mouse embryonic stem cell test (mEST) [17] and a human 3D *in vitro* model [18] were used with comparable compound concentrations. Both of the *in vitro* systems depend on the application of three-dimensional embryoid body cultures derived from either mouse embryonic stem cells (mESC, ES-D3, ATCC, CRL-1934) or human episomal induced pluripotent stem cells (hiPSC, Gibco, A18945). The teratogenicity determination of the mEST is based on the half maximal inhibition of differentiation into beating cardiomyocytes (ID_50_) and the human model focuses on the assessment of differential gene expression of early developmental markers within non-cytotoxic concentration ranges (TC_20_).

### 2.6. Compound application and experimental design

All compounds were reconstituted in sterile-filtered DMSO (Sigma-Aldrich D2438) except for penicillin, which was diluted in water and then stored at −20°C (see Table 1). The vehicle treatments used a concentration of DMSO or water equivalent to that of the highest DMSO or water concentration of the corresponding compound treatments. This ranged from 0.1-0.2% by volume DMSO and never exceeded 0.25%, and 0.8% by volume water. We used a small panel of compounds (Tab.1) which represent both positive (*all-trans* retinoic acid, valproic acid, bosentan, thalidomide, phenytoin, ibuprofen) and negative (penicillin G) references. Classifications are based on data from ICH S5 (R3) guidelines [5] (Suppl. Fig.S1). Each experiment was repeated 3 times for the RUES2-GLR and E14Tg2A cell lines and at least twice for the T/Bra::GFP line. The 96-well plate design, compound administration and timing of exposure can be seen in Figure 1. Widefield imaging data, describing both morphological shape changes over time and fluorescent reporter expression, was collected at 120 h time points in the mouse system, and at 72 h time points in the human system.

**Table 1:** Administered compounds and their respective treatment concentrations. *Phenytoin was soluble in DMSO at 200 mM but precipitates when mixed with PBS^+/+^, N2B27 or E6 medium at concentrations higher than 20 μM. Classification based on EMA ICH S5 (R3) guidelines on reproductive toxicology [5].

| Compound | Teratogenicity classification | Treatment Concentrations [X1, X2, X3] (mouse) | Treatment Concentrations [X1, X2, (X3)] (human) |
| --- | --- | --- | --- |
| All-*trans* retinoic acid | Positive | 0.4 nM, 33 nM, 100 nM | 0.4 nM, 33 nM |
| Valproic Acid | Positive | 4 μM, 333 μM, 1000 μM | 4 μM, 333 μM |
| Bosentan | Positive | 4 μM, 8 μM, 18 μM | 4 μM, 18 μM |
| Thalidomide | Positive | 0.4 μM, 11 μM, 100 μM | 0.4 μM, 100 μM |
| Phenytoin* | Positive | 1 μM, 5 μM, 20 μM | 20 μM |
| Ibuprofen | Positive | 63 μM, 120 μM, 250 μM | 63 μM, 250 μM |
| Penicillin G | Negative | 63 μM, 1 mM, 2 mM | 63 μM, 1 mM, 2 mM |

**Figure 1:**
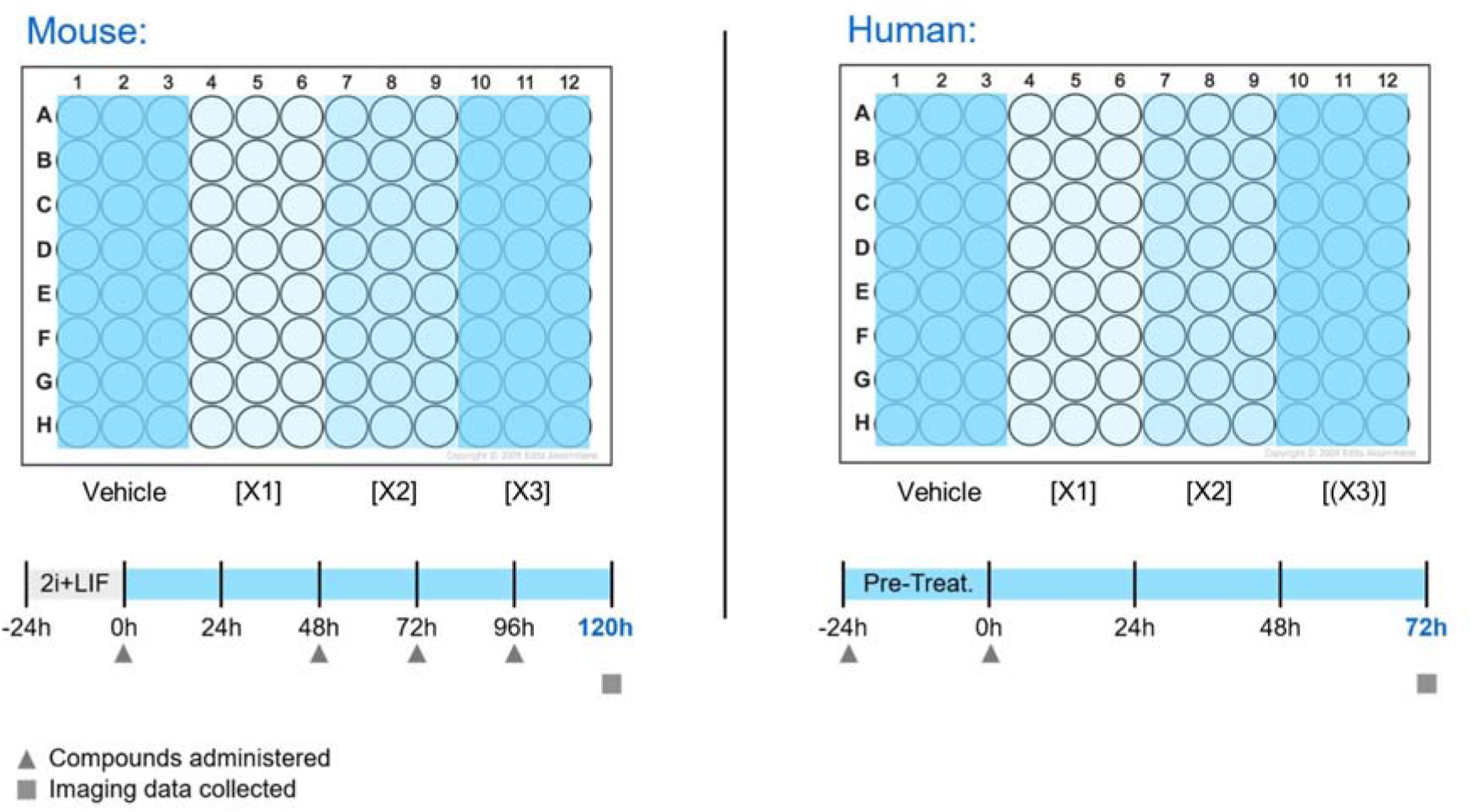
Experimental design of mouse and human gastruloid treatment. Exposure times and conditions of mouse and human gastruloid treatments. Final endpoints for study representation were obtained either after 120 h (mouse) or after 72 h (human). DMSO and water were used as vehicle controls, and X1, X2, X3 refer to the various treatment concentrations of the compounds (see Table 1).

### 2.7. Microscopy and Image Analysis

Live imaging was conducted with a Nikon Ti-E widefield microscope equipped with a cooled CMOS camera (Orca Flash 4.0, Hamamatsu) with appropriate environmental controls (37°C, 5% CO_2_; Okolab). Images were acquired through the Nikon NIS-Elements software platform. Imaging datasets were first converted to TIFF format with NIS-Elements, before further manipulation with the FIJI ImageJ distribution to collate montages per condition. Images of the mouse gastruloids were then cropped and the bit-depth of the images of both human and mouse gastruloids was reduced with custom ImageJ macros (details available on request). The 8-bit TIFF files were passed through a custom analysis pipeline, implemented in the *Project Jupyter Python 3.6* distribution [23]. Input files of the mouse gastruloids were cropped to a size that favored optimal performance of the computer vision methods used by the pipeline. Quantification contours were checked manually and the resulting measurements were curated by removing any measurements where the contours had failed to describe the shape of the human and mouse gastruloids accurately.

### 2.8. Data Analysis and Statistics

Curated datasets were handled in the *RStudio R* software environment using bespoke code and the *ggplot2, ggpubr* and *rstatix* packages. Quantification of morphology of mouse gastruloids and human gastruloids was based on contour circularity, overall area and inverse aspect ratio. The circularity measurement relates the area and perimeter of the contour to the ratio expected for a circle. The inverse aspect ratio measurement describes the proportions of the gastruloid contour by first bounding its shape within a rectangle. In all cases, the shorter side was divided by the longer side to give a number less than or equal to 1. This metric therefore measures the similarity to a regular shape: circular contours have a value of 1 while more elongated structures have increasingly small values.

Statistical comparisons were made between data from the control treatment and each experimental condition for each morphological metric. Student’s t-test was used for these comparisons with Bonferroni correction; adjusted p values are indicated with asterisks corresponding to 0 < p ≤ 1×10^-4 (****), 1×10^-4 < p ≤ 0.001 (***), 0.001 < p ≤ 0.01 (**), 0.01 < p ≤ 0.05 (*) and p > 0.05 (ns).

## 3. Results

### 3.1. All-trans-retinoic Acid

Mouse gastruloids were treated with 0.15% DMSO (vehicle control) as well as concentrations of 0.4 nM, 33 nM and 100 nM all-*trans* retinoic acid (ATRA) (Tab.1,2). Treatment with 0.4 nM was sufficient to show significant inhibition of axial extension in mouse gastruloids compared to the controls at 120 h, whereas higher dosages (33 nM, 100 nM) were associated with drastically reduced size, and gastruloids assumed a spherical shape without any axial elongation (Fig.2b,c, Fig.3a, Suppl. Fig.S3a). These observations were consistent across both lines tested, with an apparent loss of T/Bra::GFP expression in all ATRA-treated gastruloids (Fig.2c). Dosages in excess of 0.4 nM are therefore likely to be cytotoxic or cytostatic in the mouse system.

**Table 2.** Summary of observed effects by compound concentration. Different concentrations of compounds with teratogenic and non-teratogenic profiles, treated on mouse and human gastruloids. Darker cell colours indicate more pronounced effects.

|  | **Mouse Gastruloids** |  |  | **Human Gastruloids** |  |  |
| --- | --- | --- | --- | --- | --- | --- |
|  | **0.4 nM** | **33 nM** | **100 nM** | **0.4 nM** | **33 nM** |  |
| **Retinoic Acid** | Morphological effect, Gene expression effect | Morphological effect; Size reduction; Suspected cytotoxicity, Gene expression effect | Morphological effect; Size reduction; Suspected cytotoxicity, Gene expression effect | Small morphological effect | Morphological effect; Size reduction, Gene expression effect |  |
|  | **4 μM** | **333 μM** | **1 mM** | **4 μM** | **333 μM** |  |
| **Valproic Acid** | Small morphological effect | Morphological effect; Size reduction; Suspected cytotoxicity, Gene expression effect | Morphological effect; Size reduction; Suspected cytotoxicity, Gene expression effect | Minimal effect | Morphological effect; Size reduction, Gene expression effect |  |
|  | **4 μM** | **8 μM** | **18 μM** | **4 μM** | **18 μM** |  |
| **Bosentan** | Minimal effect | Minimal effect | Minimal effect | Minimal effect | Morphological effect; Size reduction, Gene expression effect |  |
|  | **0.4 μM** | **11 μM** | **100 μM** | **0.4 μM** | **100 μM** |  |
| **Thalidomide** | No effect | No effect | Gene expression effect | Morphological effect; Size reduction, Gene expression effect | Morphological effect; Size reduction, Gene expression effect |  |
|  | **1 μM** | **5 μM** | **20 μM** | **20 μM** | **-** |  |
| **Phenytoin** | No effect | Gene expression effect | Gene expression effect | Minimal effect | **-** |  |
|  | **63 μM** | **120 μM** | **250 μM** | **63 μM** | **250 μM** |  |
| **Ibuprofen** | Gene expression effect | Gene expression effect | Gene expression effect | Morphological effect; Size reduction, Gene expression effect | Morphological effect; Size reduction, Gene expression effect |  |
|  | **63 μM** | **1 mM** | **2 mM** | **63 μM** | **1 mM** | **2 mM** |
| **Penicillin G** | No effect | No effect | No effect | No effect | No effect | No effect |

**Figure 2:**
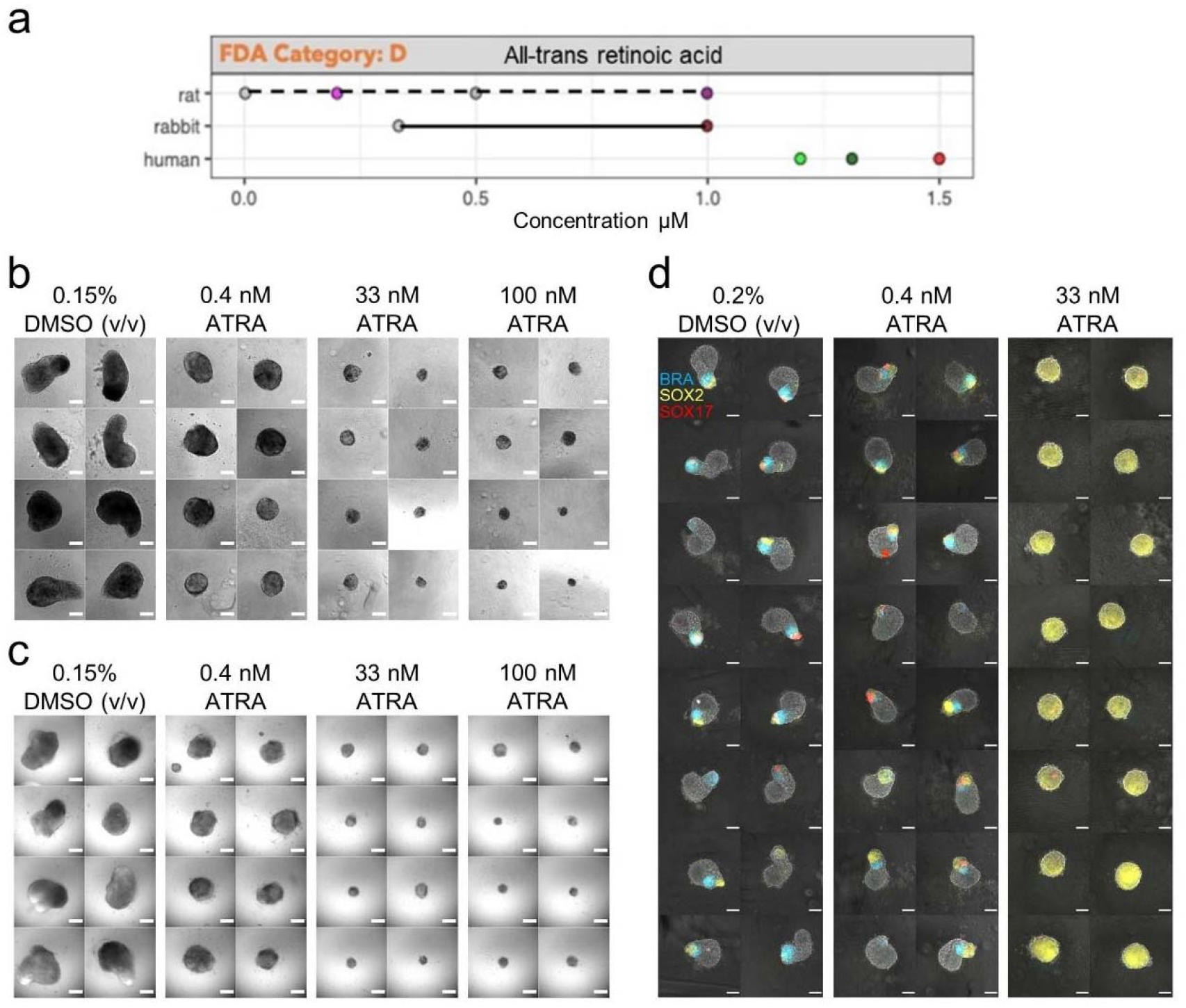
Gastruloids following *all-trans* retinoic acid exposure. (a) Literature-based exposure limits in different species in [μM] (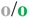 no effect/ NOAEL; 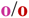 teratogenic/ LOAEL, see Suppl. Fig.S1). (b-c) E14Tg2A (b) and T/Bra::GFP (c) mouse gastruloids at 120h, following exposure to DMSO (vehicle control) or *all-trans* retinoic acid (ATRA). (d) RUES2-GLR human gastruloids at 72h, following exposure to DMSO or ATRA. Color indicates fluorescent expression of BRA-mCerulean (blue), SOX2-mCitrine (yellow) and SOX17-tdTomato (red). Scale bars represent 200 μm (b,c) and 100μm (d).

**Figure 3:**
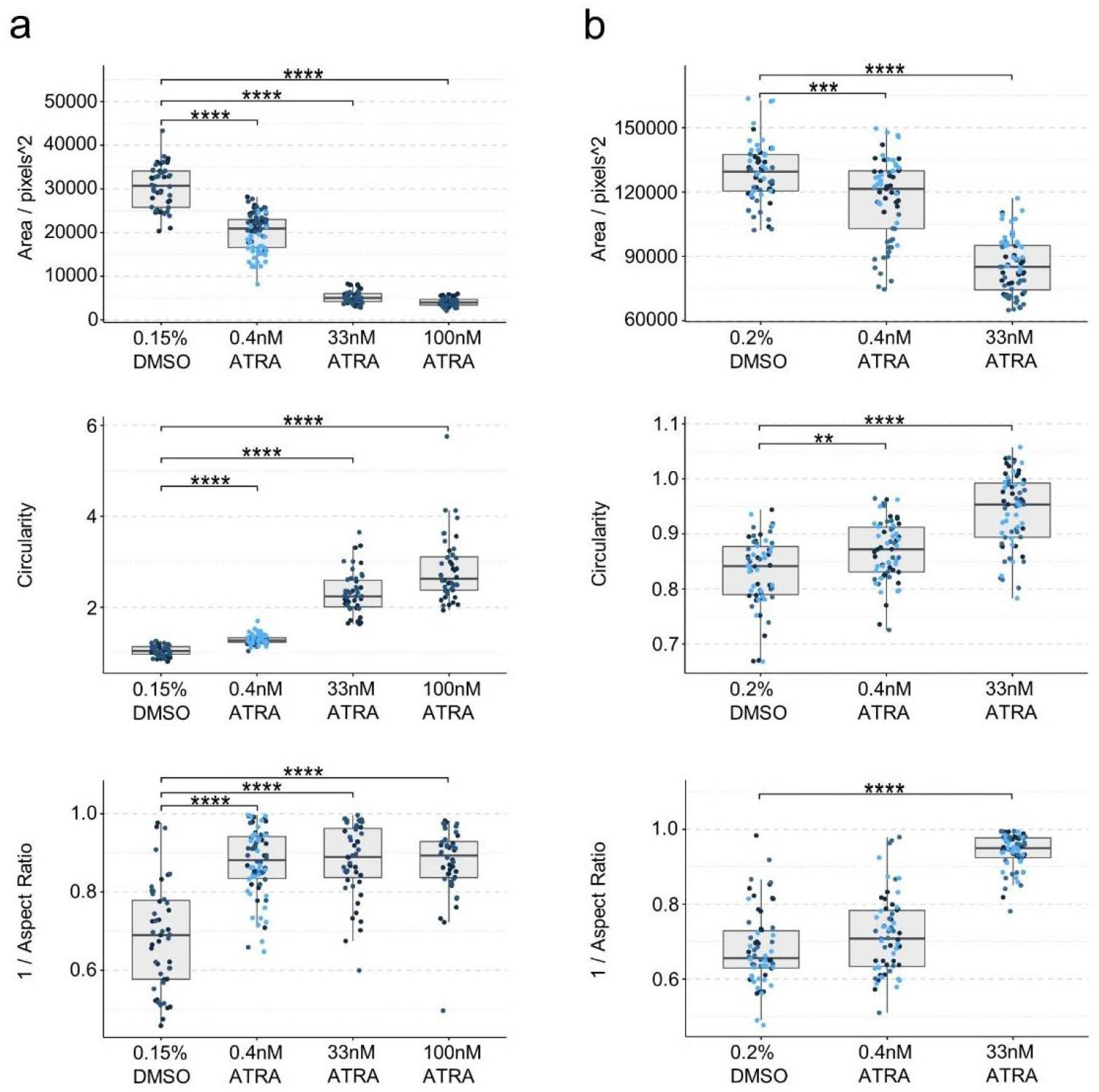
Gastruloid quantification following *all-trans* retinoic acid exposure. Quantification of morphology of mouse gastruloids (a), and human gastruloids (b) including Area (top), Circularity (middle) and 1/Aspect Ratio (bottom). Dot colors indicate experimental replicates, and boxplots indicate spread of the data. Significant differences between DMSO (vehicle control) and *all-trans* retinoic acid (ATRA) treatment conditions are indicated in the plots (Student’s t-test, Bonferroni corrected. Adjusted p-values less than 0.05 are indicated by asterisks (see Materials and Methods for thresholds)).

In contrast to these observations, human gastruloids treated with 0.4 nM retinoic acid appeared to maintain an equivalent representation of the three germ layers and underwent partial axial elongation (Fig.2d), although they were still significantly smaller and more circular than controls at 72 h (Fig.3b). Concentrations of 33 nM retinoic acid completely inhibited axial extension and promoted the over-expression of *SOX2*, as observed by fluorescent signal (Fig.2d). There was no detectable *BRA* expression across the three replicates. We also observed that, on average, 42% of human gastruloids treated with 33 nM retinoic acid expressed *SOX17* at low levels, compared to 88% of DMSO controls, where expression was higher. Taken together, these results suggest a lowest observable adverse effect level (LOAEL) of 0.4 nM for both mouse and human gastruloids, albeit with slightly different observable effects.

In comparison, maximum plasma concentrations of the lowest observed adverse effect level (LOAEL Cmax) in rats and rabbits range around 1 μM and human therapeutic plasma concentration (human Cmax) is about 1.3 μM (Fig.2a, Tab.3) [5]. We therefore compared our data with *in vitro* studies of mouse and human embryoid bodies (EB), which revealed ID_50_ concentrations (half maximal inhibitory concentration of cardiomyocyte differentiation) ranging between 3 nM (mEST) and TC_20_ concentrations (threshold concentration of 20% teratogenicity induction based on differential gene expression) of 0.1 nM in hiPSC-derived EBs for ATRA, which corresponds well to the range of the gastruloid assay (Tab.3, Suppl. Fig.S2).

**Table 3:** Effective concentrations. The table shows the lowest observable adverse effect level (LOAEL) concentrations for mouse and human gastruloids in comparison with rat *in vivo* Cmax LOAEL data, ID_50_ values (half maximal inhibitory concentration of cardiomyocyte differentiation) detected by mEST or TC_20_ (threshold concentration where 20% of teratogenicity was detected by differential expression) from hiPSC-derived EBs, and the maximum human therapeutic plasma concentrations (Cmax) [5].

|  | Mouse |  |  | Human |  |  |
| --- | --- | --- | --- | --- | --- | --- |
| Compound | mEST ID<sub>50</sub> | mouse Gastruloid LOAEL | rat *in vivo* LOAEL Cmax | hiPSC-derived EBs TC<sub>20</sub> | Human Gastruloid LOAEL | human *in vivo* Cmax |
| Retinoic Acid | 3.7 nM | 0.4 nM | 1 µM | < 0.1 nM | 0.4 nM | 1.31 µM |
| Valproic Acid | 589 µM | 333 µM | 1.6 mM | 85 µM | 333 µM | 1.4 mM |
| Bosentan | 68.2 µM | n/a | 30 µM | 13 µM | 18 µM | n/a |
| Thalidomide | n/a | 100 µM | 19 µM | < 0.1 µM | 0.4 µM | 2.4 µM |
| Phenytoin | 0.39 µM | 5 µM | 106 µM | < 8 µM | n/a | 57 µM |
| Ibuprofen | 49 µM | 63µM | 1.6 mM | 855 µM | 63 µM | 286 µM |
| Penicillin G | n/a | n/a | n/a | n/a | n/a | n/a |

### 3.2. Valproic Acid

Treatments with 4 μM valproic acid seemed to be fairly well tolerated in both the mouse and human gastruloid systems, albeit with an increase in circularity in the mouse, and a decrease in size in the human (Fig.5a-b). Concentrations higher than 333 μM produced clearly spherical mouse gastruloids (Fig.4b) of reduced size (Fig.5a), indicating a loss of axial elongation morphogenesis with impaired cell viability or proliferation at 1 mM exposure compared to DMSO controls. These observations were replicated in the T/Bra::GFP cell line, which maintained localised GFP expression after treatment with 4 μM valproic acid and did not show significantly increased circularity (Fig.4c, Suppl. Fig.S3b). Higher concentrations resulted in smaller gastruloids, impaired elongation and a loss of GFP expression.

**Figure 4:**
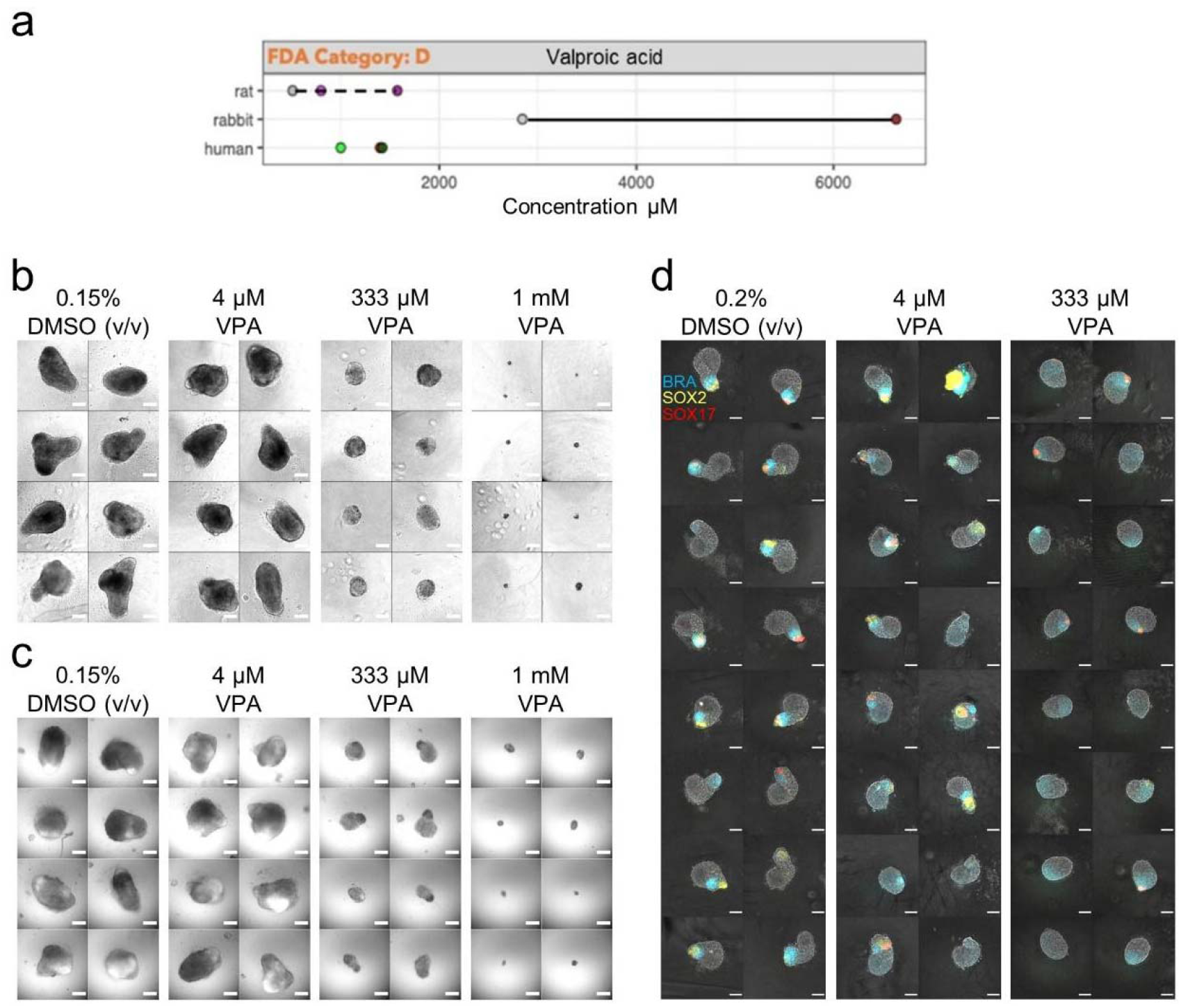
Gastruloids following valproic acid exposure. (a) Literature-based exposure limits in different species in [μM] (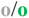 no effect/ NOAEL; 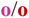 teratogenic/ LOAEL, see Suppl. Fig.S1). (b-c) E14Tg2A (b) and T/Bra::GFP (c) mouse gastruloids at 120h, following exposure to DMSO (vehicle control) or valproic acid (VPA). (d) RUES2-GLR human gastruloids at 72h, following exposure to DMSO or valproic acid. Color indicates fluorescent expression of BRA-mCerulean (blue), SOX2-mCitrine (yellow) and SOX17-tdTomato (red). Scale bars represent 200 μm (b,c) and 100μm (d).

**Figure 5:**
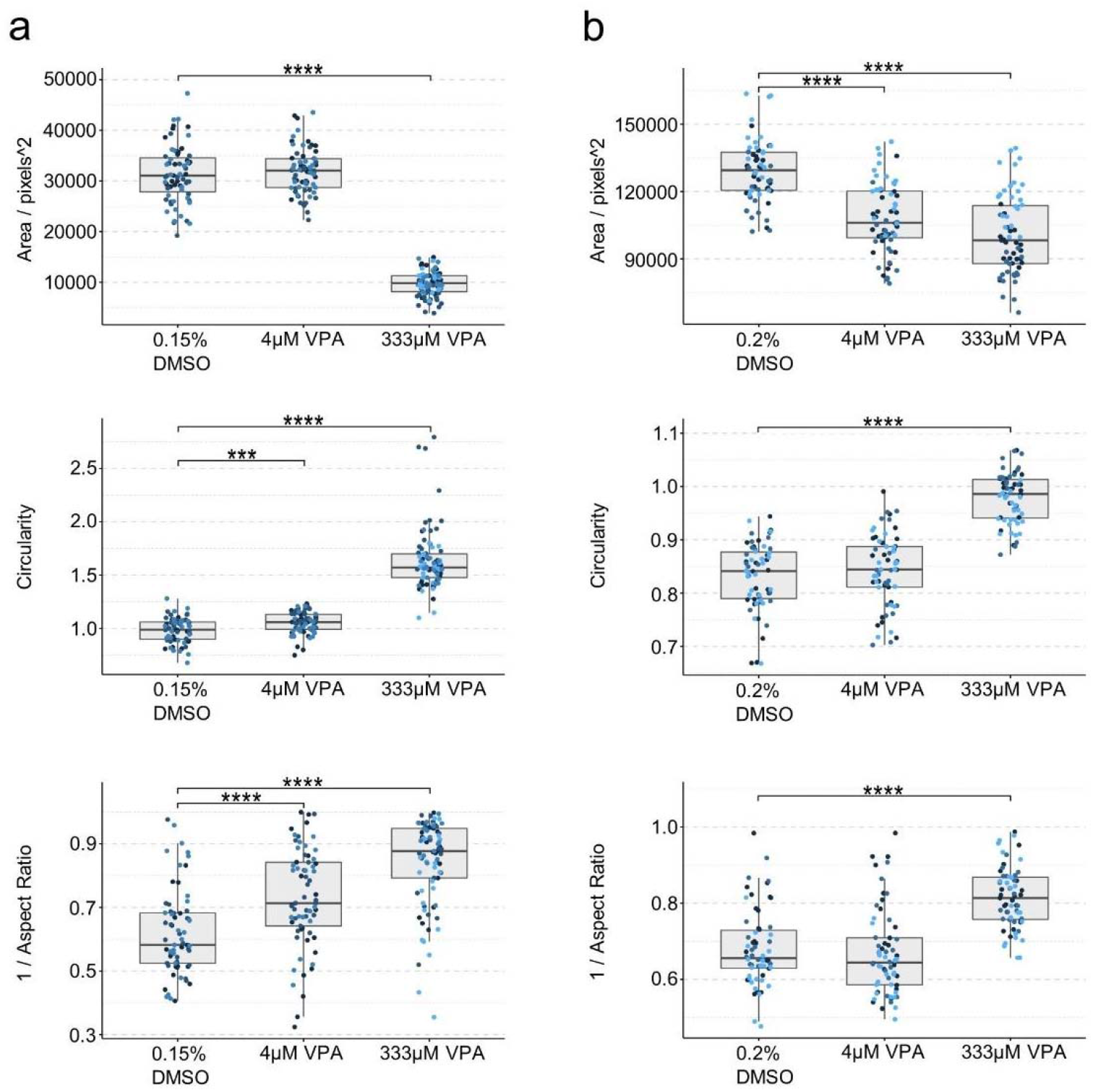
Gastruloid quantification following valproic acid exposure. Quantification of morphology of mouse gastruloids (a), and human gastruloids (b) including Area (top), Circularity (middle) and 1/Aspect Ratio (bottom). Dot colors indicate experimental replicates, and boxplots indicate spread of the data. Significant differences between DMSO (vehicle control) and valproic acid (VPA) treatment conditions are indicated in the plots (Student’s t-test, Bonferroni corrected. Adjusted p-values less than 0.05 are indicated by asterisks (see Materials and Methods for thresholds)).

Human gastruloids treated with 333 μM valproic acid also failed to undergo axial elongation morphogenesis. On examining reporter expression, these gastruloids were found to show very low *SOX2* expression and elevated *BRA* expression in comparison to 0.2% DMSO (v/v) controls (Fig.4d, Fig.5b). These results indicate a common LOAEL of 333 μM for valproic acid (VPA) for both the mouse and human gastruloid systems. Compared to this, VPA studies in the mEST revealed an ID_50_ of 589 μM for the inhibition of D3 mouse cardiomyocytes, whereas studies in human iPSC derived EBs revealed a TC_20_ of 85 μM (Tab.3, Suppl. Fig.S2). By contrast, data for *in vivo* LOAEL Cmax are 1.6 mM in rat and 6.6 mM in rabbits, whereas human Cmax values are about 1.4 mM (Fig.4a, Tab.3) [5].

### 3.3. Bosentan

Mouse gastruloids treated with 4 to 18 μM bosentan were slightly larger than controls, but did not appear to be defective in morphogenesis (Fig.6b, Fig.7a). Gastruloids generated from the T/Bra::GFP cell line cultured in the presence of bosentan showed robust GFP expression that was localised to the pole of the elongations, as observed in untreated controls (Fig.6c). Morphologically, they were larger and more elongated than controls with 4 μM and 8 μM treatments (Suppl. Fig.S3c). Bosentan was predicted as a false negative with the mEST. Further investigation of the mouse system will be required to determine whether broader patterns of gene expression are dysregulated under comparable culture conditions. Similarly, human gastruloids treated with 4 μM bosentan appeared to maintain representation of all three germ layers and axial elongation morphogenesis, although they were marginally smaller than DMSO-treated gastruloids (Fig.6d, Fig.7b).

**Figure 6:**
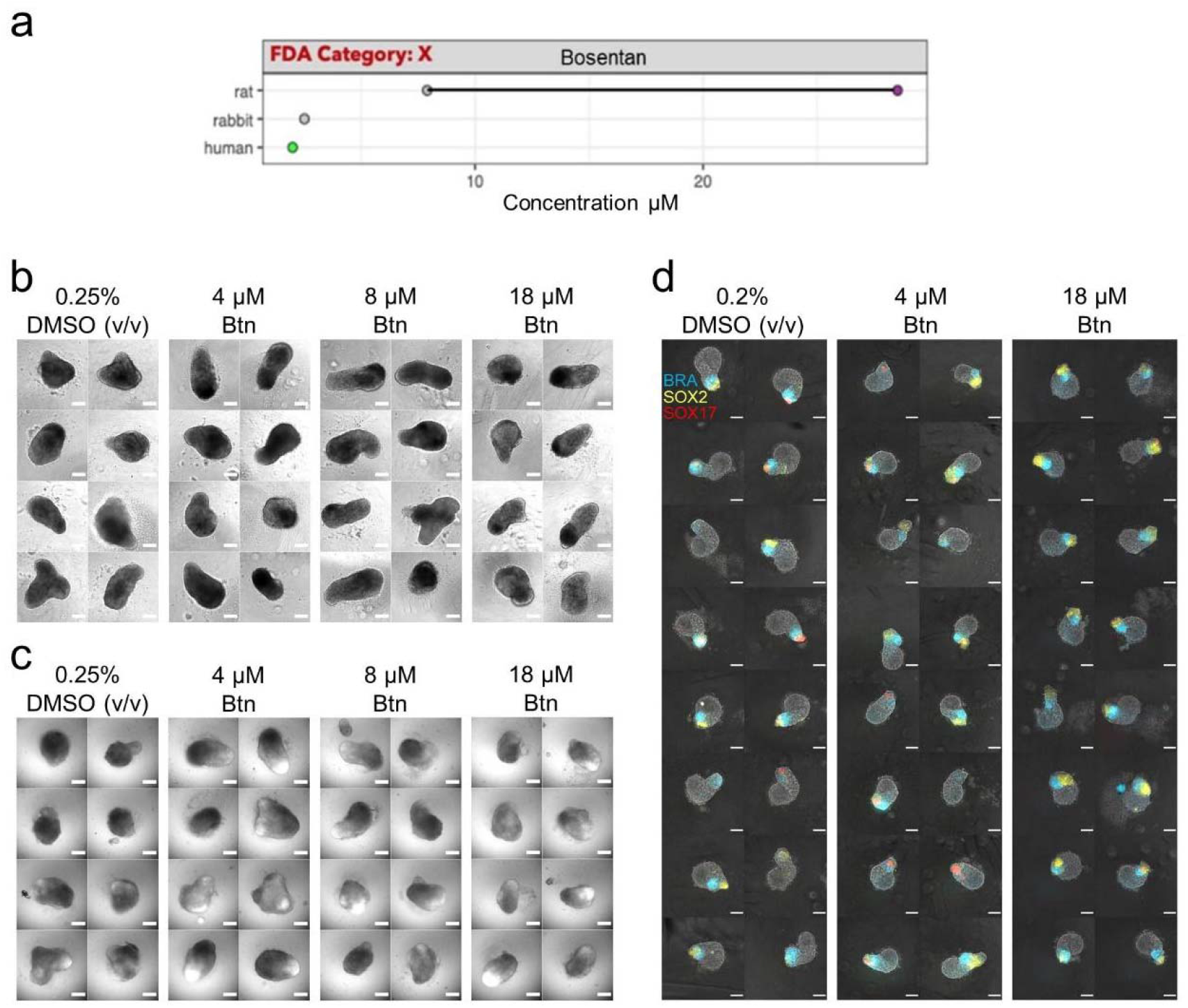
Bosentan exposure. (a) Literature-based exposure limits in different species in [μM] (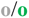 no effect/ NOAEL; 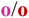 teratogenic/ LOAEL, see Suppl. Fig.S1). (b-c) E14Tg2A (b) and T/Bra::GFP (c) mouse gastruloids at 120h, following exposure to DMSO (vehicle control) or bosentan (Btn). (c) RUES2-GLR human gastruloids at 72h, following exposure to DMSO or bosentan. Color indicates fluorescent expression of BRA-mCerulean (blue), SOX2-mCitrine (yellow) and SOX17-tdTomato (red). Scale bars represent 200 μm (b,c) and 100μm (d).

**Figure 7:**
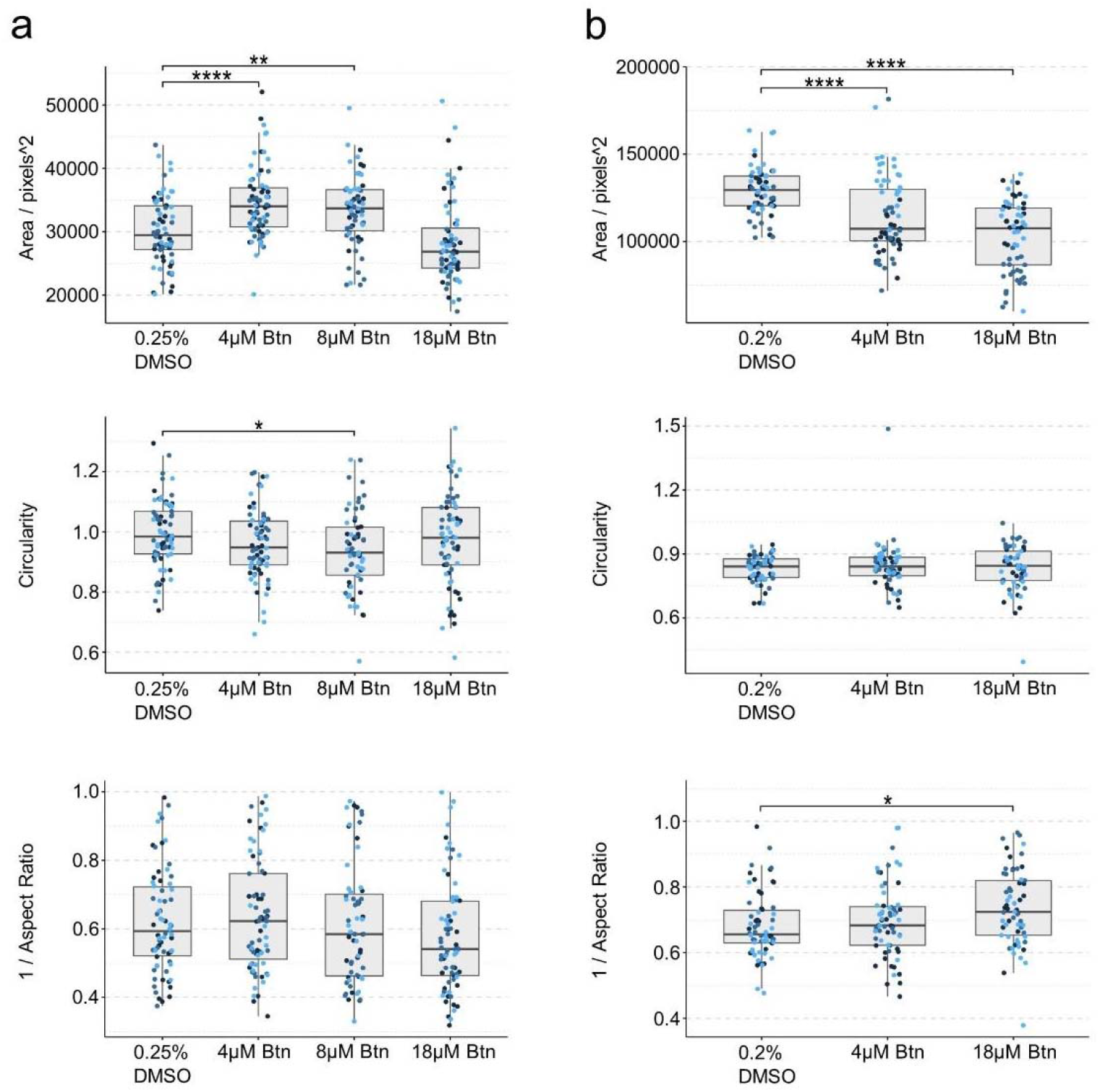
Gastruloid quantification following bosentan exposure. Quantification of morphology of mouse gastruloids (a), and human gastruloids (b) including Area (top), Circularity (middle) and 1/Aspect Ratio (bottom). Dot colors indicate experimental replicates, and boxplots indicate spread of the data. Significant differences between DMSO (vehicle control) and bosentan (Btn) treatment conditions are indicated in the plots (Student’s t-test, Bonferroni corrected. Adjusted p-values less than 0.05 are indicated by asterisks (see Materials and Methods for thresholds)).

At 18 μM, however, the human gastruloids appeared to lose expression of *SOX17*. We observed that, on average, 33% showed low levels of *SOX17* expression in comparison to 88% of vehicle controls (0.2% DMSO (v/v)), where expression was higher (Fig.6d, Suppl. Fig.S4a). The results from the human gastruloid cultures therefore suggest a LOAEL of 18 μM bosentan. This corresponds well with the TC_20_ value for hiPSC-derived embryoid bodies that was determined at 13 μM. The *in vivo* LOAEL Cmax for bosentan is about 30 μM in rats and it showed no effects in rabbits (Fig.6a, Tab.3) [5].

### 3.4. Thalidomide

Mouse gastruloids treated with 0.4-100 μM ±-thalidomide were not significantly altered in either size or morphology (Fig.8b,c, Fig.9a, Suppl. Fig.S4b) for both lines tested. Increasing concentrations appeared to correlate with reduced GFP expression in the T/Bra::GFP cell line (100 μM). Thalidomide is false negative predicted with the mEST at concentrations up to 2 mM and did not show any inhibition of cardiomyocytes. Human gastruloids, on the other hand, showed clearly increased *SOX2* expression following exposure to both 0.4 μM and 100 μM ±- thalidomide (Fig.8d). This was accompanied by significant decreases in gastruloid size (Fig.9b). These results suggest a greater sensitivity of the human gastruloid system to exposure to ±- thalidomide, with a LOAEL for this system defined at 0.4 μM in human gastruloids and no observable morphological effect in mouse gastruloids at the concentrations studied. Investigations made with the human iPSC derived EB model revealed differential gene expression effects at TC_2_0 concentrations < 0.1 μM compared to DMSO vehicle controls (Tab.3, Suppl. Fig.S2). *In vivo* studies revealed a LOAEL Cmax of 19 μM in rats and 8.4 μM in rabbits [5]. Human therapeutic plasma concentrations are about 2.4 μM (Fig.8a, Tab.3). These results therefore suggest that gastruloids could be used to identify species-specific responses to chemical exposure.

**Figure 8:**
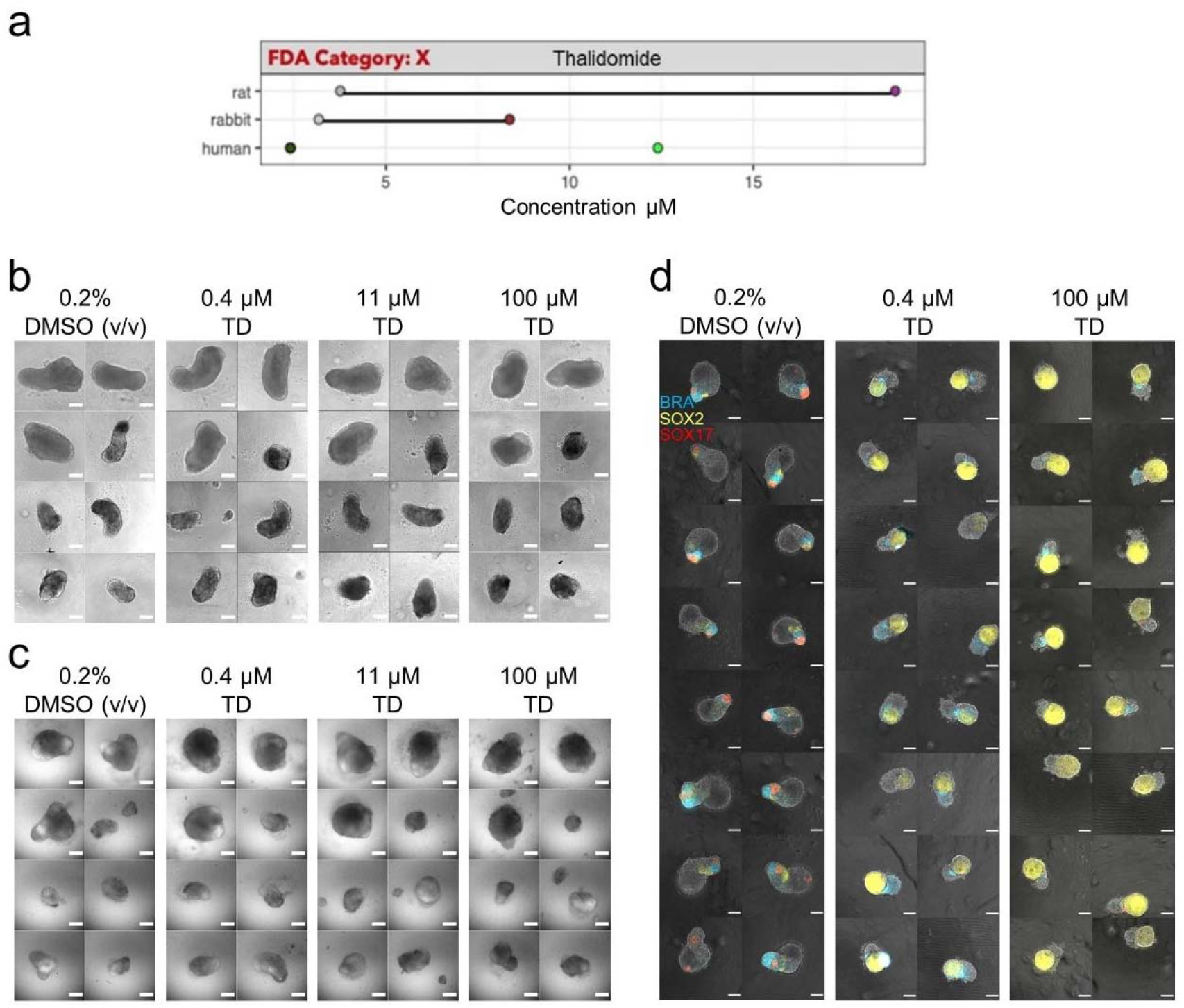
Thalidomide Exposure. (a) Literature-based exposure limits in different species in [μM] (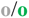 no effect/ NOAEL; 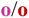 teratogenic/ LOAEL, see Suppl. Fig.S1). (b-c) E14Tg2A (b) and T/Bra::GFP (c) mouse gastruloids at 120h, following exposure to DMSO (vehicle control) or thalidomide (TD). (d) RUES2-GLR human gastruloids at 72h, following exposure to DMSO or thalidomide. Color indicates fluorescent expression of BRA-mCerulean (blue), SOX2-mCitrine (yellow) and SOX17-tdTomato (red). Scale bars represent 200 μm (b,c) and 100μm (d).

**Figure 9:**
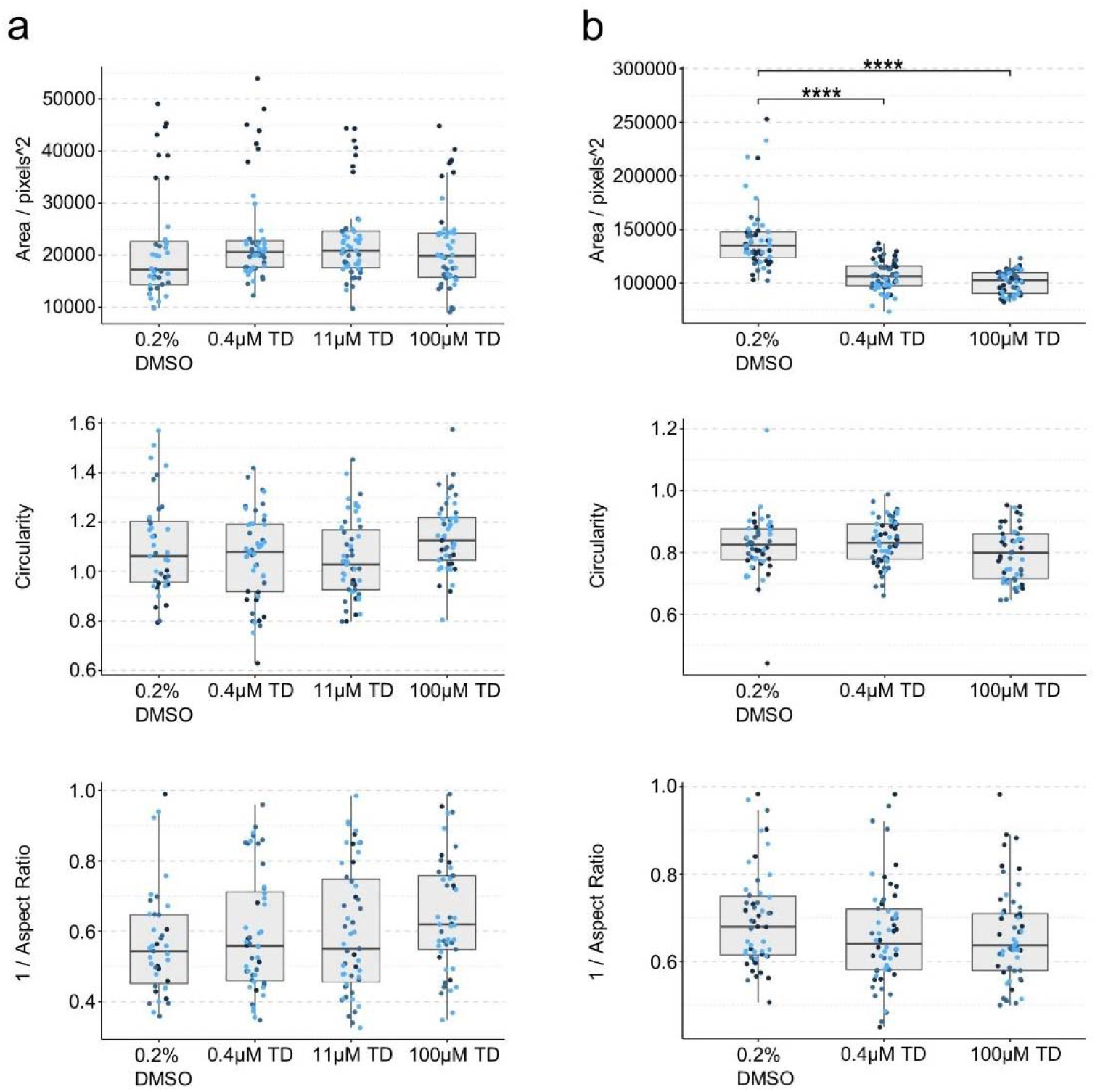
Gastruloid quantification following thalidomide exposure. Quantification of morphology of mouse gastruloids (a), and human gastruloids (b) including Area (top), Circularity (middle) and 1/Aspect Ratio (bottom). Dot colors indicate experimental replicates, and boxplots indicate spread of the data. Significant differences between DMSO (vehicle control) and thalidomide (TD) treatment conditions are indicated in the plots (Student’s t-test, Bonferroni corrected. Adjusted p-values less than 0.05 are indicated by asterisks (see Materials and Methods for thresholds)).

### 3.5. Phenytoin

Mouse gastruloids treated with 1-20 μM phenytoin were not significantly different compared to the vehicle-only controls in terms of size and morphology (0.1% DMSO (v/v); Fig. 10b,c, Fig. 11a, Suppl. Fig.S4c). The 20 μM treatment in the T/Bra::GFP line was the only exception, which was significantly smaller in these data. The expression of T/Bra::GFP appeared to be negatively correlated with increasing concentrations of phenytoin (> 5μM, Fig. 10c). Human gastruloids treated with 1-20 μM phenytoin maintained proportions and patterns of gene expression that were comparable to controls (Fig.10d, Fig. 11b). It would be reasonable to conclude that the 1-20 μM concentrations tested fall within the NOAEL (no observable adverse effect level) for these systems. Any follow-up study would be well-advised to use this compound in a different solvent that could allow higher concentrations of phenytoin to be used experimentally without precipitation. Particularly, inhibition of cardiomyocyte-differentiation has already been detected with the mEST at ID_50_ concentrations of 49 μM, while with the hiPSC-derived EB model we observed differential gene expression effects at TC_20_ concentrations of < 8 μM for phenytoin. *In vivo* studies revealed a LOAEL Cmax of about 106 μM in rats and 135 μM in rabbits [5]. Human therapeutic Cmax is about 57 μM (Fig.10a, Tab.3).

**Figure 10:**
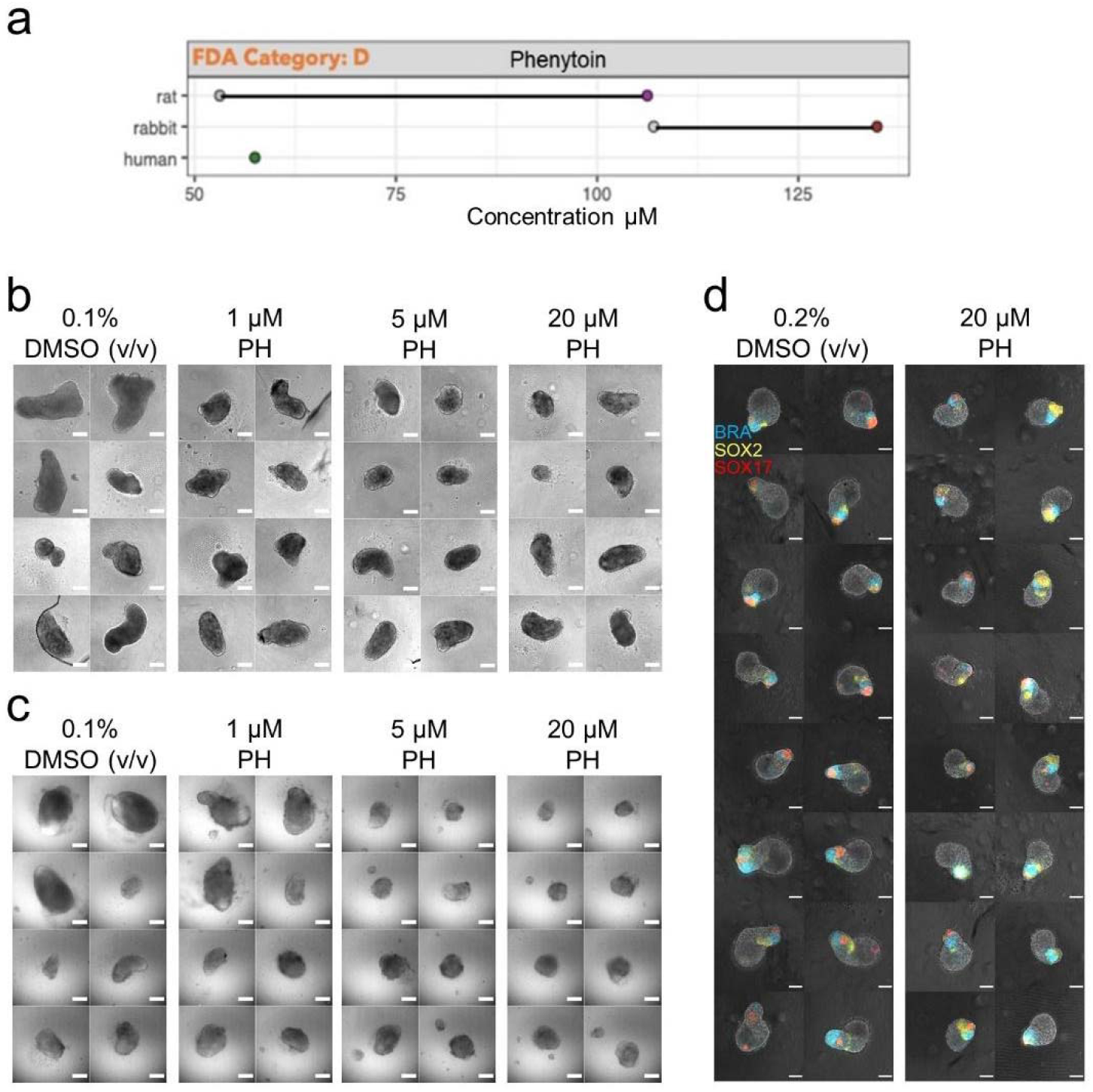
Phenytoin Exposure. (a) Literature-based exposure limits in different species in [μM] (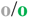 no effect/ NOAEL; 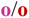 teratogenic/ LOAEL, see Suppl. Fig.S1). (b-c) E14Tg2A (b) and T/Bra::GFP (c) mouse gastruloids at 120h, following exposure to DMSO (vehicle control) or phenytoin (PH). (d) RUES2-GLR human gastruloids at 72h, following exposure to DMSO or phenytoin. Color indicates fluorescent expression of BRA-mCerulean (blue), SOX2-mCitrine (yellow) and SOX17-tdTomato (red). Scale bars represent 200 μm (b,c) and 100μm (d).

**Figure 11:**
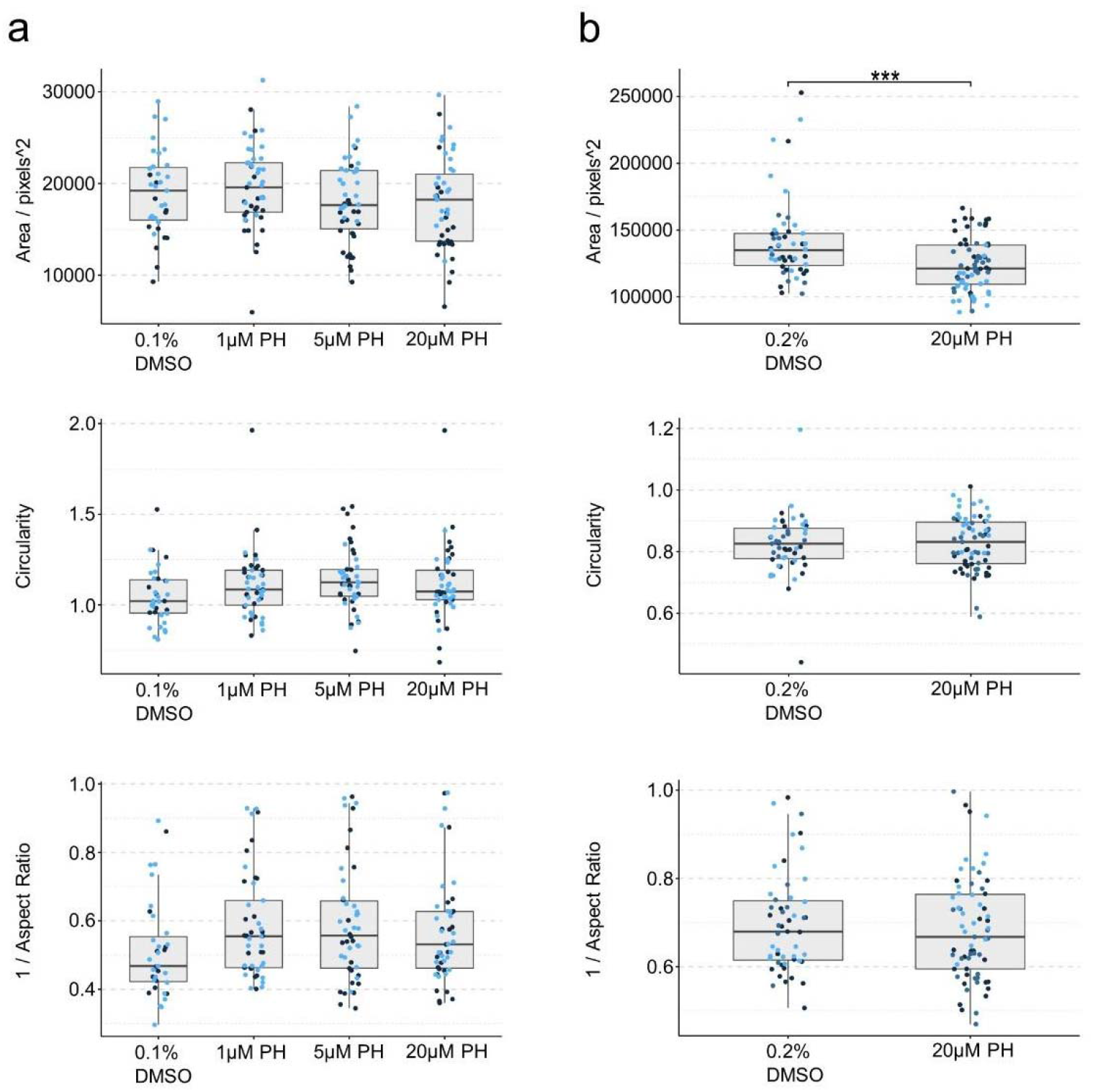
Gastruloid quantification following phenytoin exposure. Quantification of morphology of mouse gastruloids (a), and human gastruloids (b) including Area (top), Circularity (middle) and 1/Aspect Ratio (bottom). Dot colors indicate experimental replicates, and boxplots indicate spread of the data. Significant differences between DMSO (vehicle control) and phenytoin (PH) treatment conditions are indicated in the plots (Student’s t-test, Bonferroni corrected. Adjusted p-values less than 0.05 are indicated by asterisks (see Materials and Methods for thresholds)).

### 3.6. Ibuprofen

Mouse gastruloids treated with 63-250 μM ibuprofen showed no consistent effect on growth or morphology in comparison to vehicle-only controls (0.12% DMSO (v/v); Fig.12b,c, Fig.13a, Suppl. Fig.S5a). The 63 μM treatment resulted in significantly less circular gastruloids, while the 120 μM treatment resulted in significantly smaller gastruloids. The T/Bra::GFP reporter, however, showed a marked increase in expression with all ibuprofen treatments tested, indicating dysregulated gene expression in comparison to controls (Fig. 12c). Preliminary experiments showed that treatments in the range of 500 μM-2 mM were cytotoxic to both mouse and human embryonic stem cells (data not shown).

**Figure 12:**
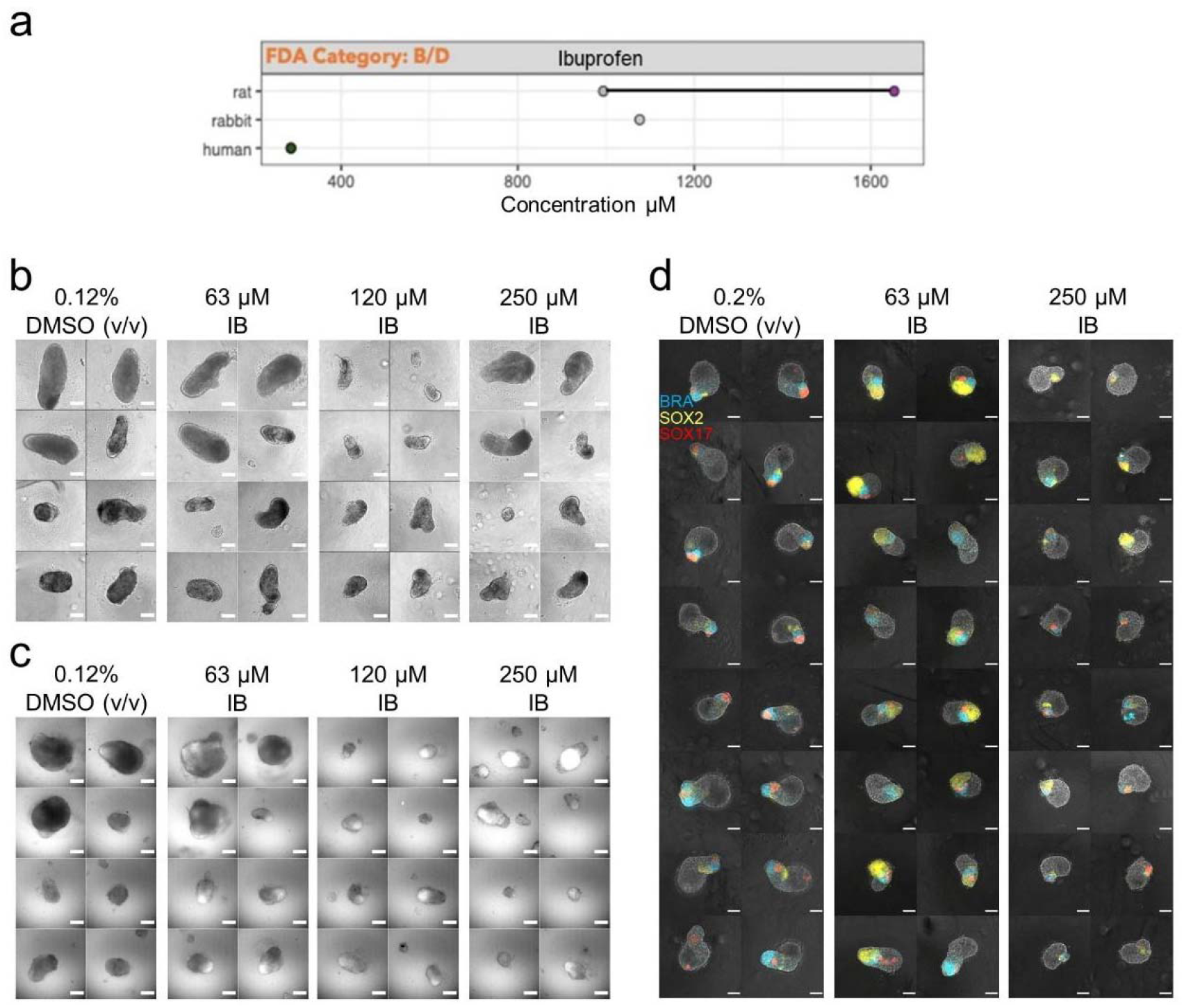
Ibuprofen Exposure. (a) Literature-based exposure limits in different species in [μM] (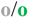 no effect/ NOAEL; 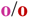 teratogenic/ LOAEL, see Suppl. Fig.S1). (b-c) E14Tg2A (b) and T/Bra::GFP (c) mouse gastruloids at 120h, following exposure to DMSO (vehicle control) or ibuprofen (IB). (d) RUES2-GLR human gastruloids at 72h, following exposure to DMSO or ibuprofen. Color indicates fluorescent expression of BRA-mCerulean (blue), SOX2-mCitrine (yellow) and SOX17-tdTomato (red). Scale bars represent 200 μm (b,c) and 100μm (d).

**Figure 13:**
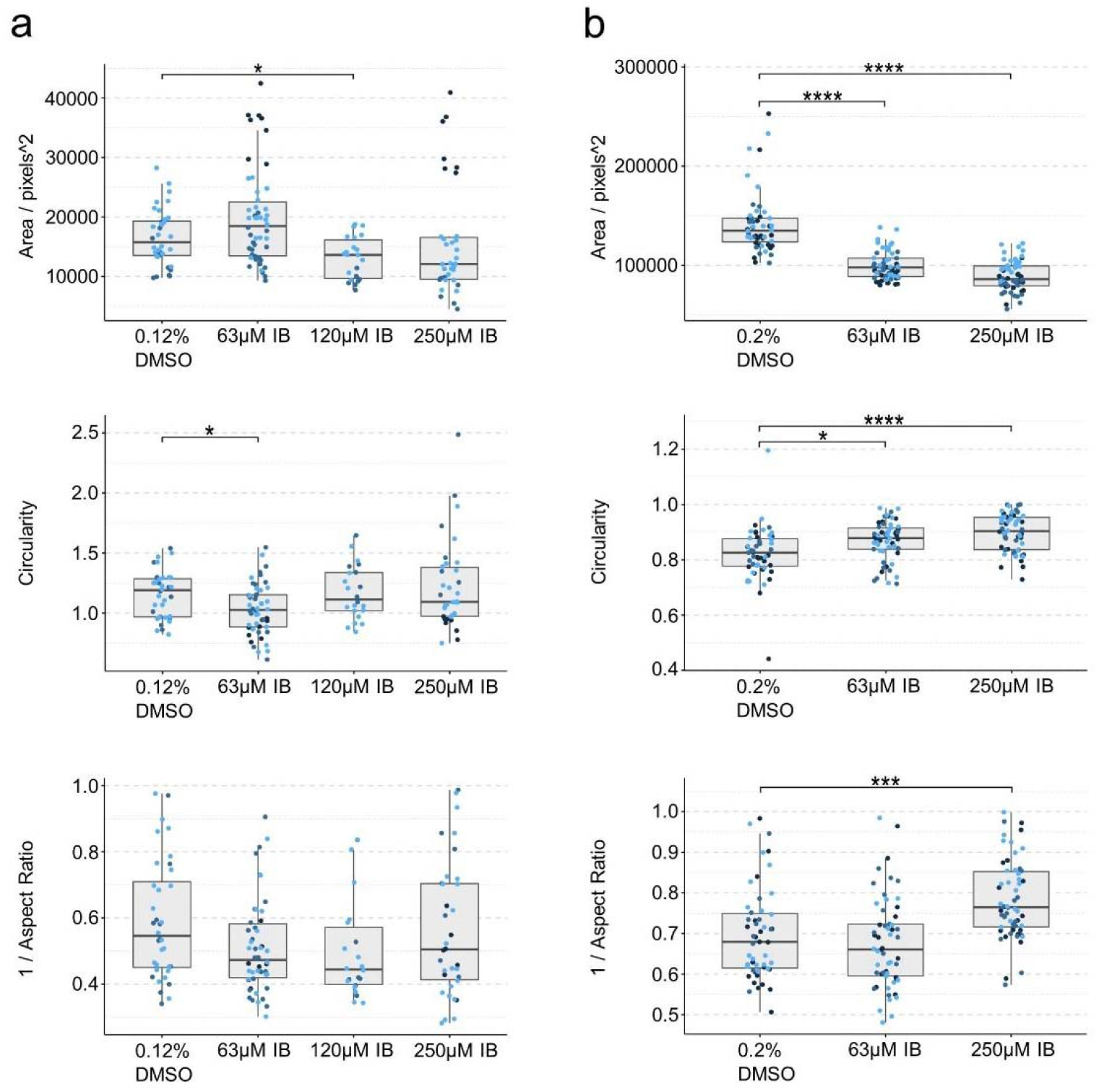
Gastruloid quantification following ibuprofen exposure. Quantification of morphology of mouse gastruloids (a), and human gastruloids (b) including Area (top), Circularity (middle) and 1/Aspect Ratio (bottom). Dot colors indicate experimental replicates, and boxplots indicate spread of the data. Significant differences between DMSO (vehicle control) and ibuprofen (IB) treatment conditions are indicated in the plots (Student’s t-test, Bonferroni corrected. Adjusted p-values less than 0.05 are indicated by asterisks (see Materials and Methods for thresholds)).

On culturing human gastruloids in the presence of 63 μM ibuprofen, the expression of *SOX2* seemed to be marginally elevated compared to vehicle-only controls (0.2% DMSO (v/v); Fig.12d). Exposure at both 63 μM and 250 μM produced gastruloids of reduced size and increased circularity (Fig. 13b). It would therefore seem that ibuprofen has a potential effect on expression of *T/Bra* in mouse and *SOX2* in human gastruloids, with a LOAEL at 63 μM under these conditions. The mEST predicted ibuprofen as negative whereas the hiPSC-derived EB model determined ibuprofen as positive substance at much higher concentrations, with a TC_20_ of 855 μM (Tab.3, Suppl. Fig.S2). The LOAEL Cmax for ibuprofen in rats is about 1.6 mM and no effects have been observed in rabbits [5]. Human therapeutic Cmax is about 286 μM (Fig.12a, Tab.3).

### 3.7. Penicillin G

Penicillin was the only negative reference compound tested beside the DMSO or water vehicle controls to identify non-specific effects. Penicillin was tested at concentrations of 63 μM, 1 mM and 2 mM (diluted in water) in the mouse and the human gastruloid model (Tab.1). No observable adverse effects on morphology or reporter gene expression were detected up to the highest concentration levels in both systems (Fig.14b-c, Fig.15a-b, Suppl. Fig.S5b). The only significant morphological changes resulted in larger, more elongated or less circular gastruloids compared to the controls. These results are consistent with other *in vitro* and *in vivo* systems as well as with human data, where penicillin is not teratogenic within therapeutic concentration levels (Fig.14a) [5].

**Figure 14:**
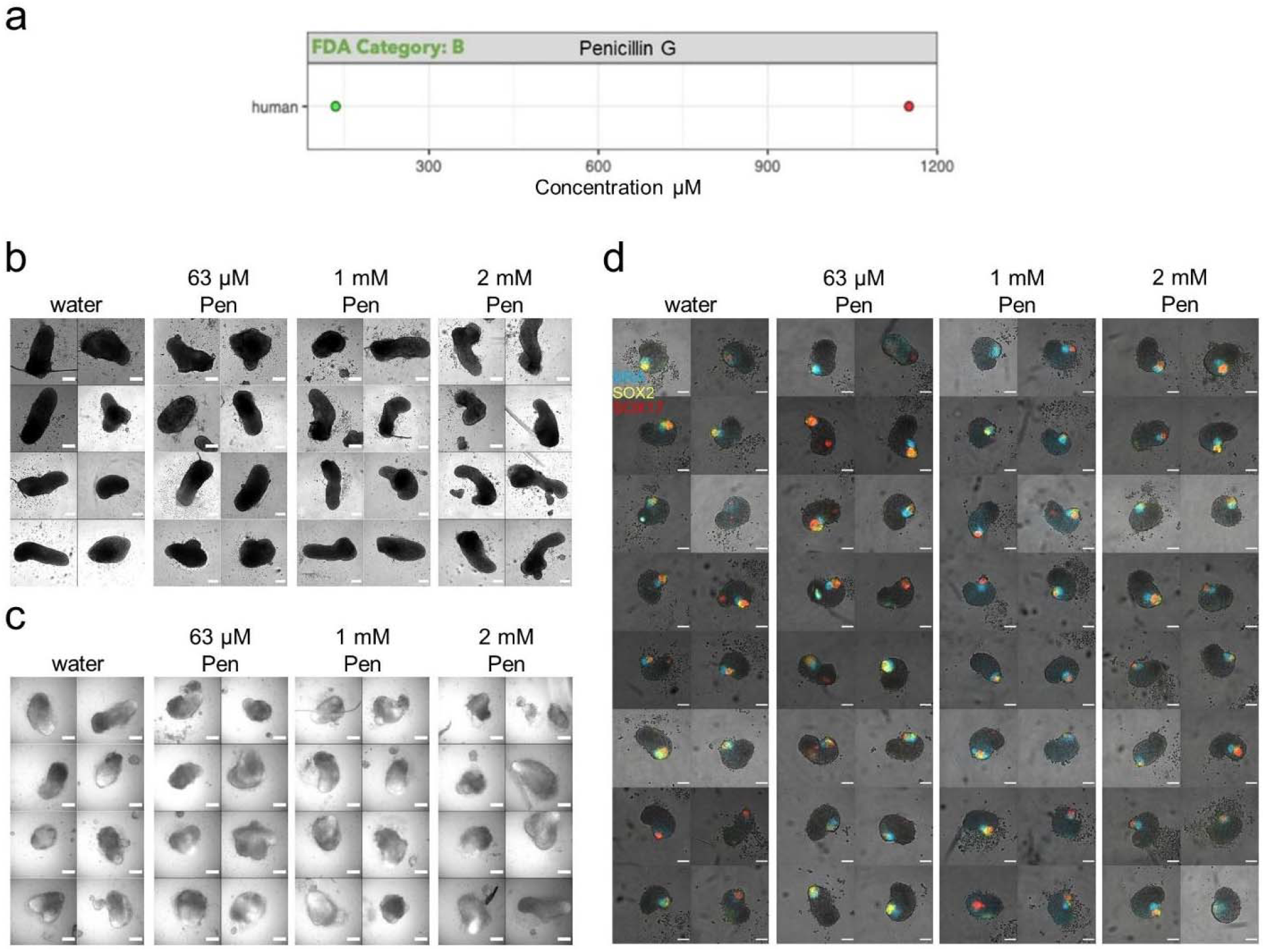
Penicillin G Exposure. (a) Literature-based exposure limits in different species in [μM] (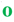 no effect/ NOAEL; 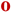 teratogenic/ LOAEL, see Suppl. Fig.S1). (b-c) E14Tg2A (b) and T/Bra::GFP (c) mouse gastruloids at 120h, following exposure to water (vehicle control) or penicillin G (Pen). (d) RUES2-GLR human gastruloids at 72h, following exposure to water or penicillin G. Color indicates fluorescent expression of BRA-mCerulean (blue), SOX2-mCitrine (yellow) and SOX17-tdTomato (red). Scale bars represent 200 μm (b,c) and 100μm (d).

**Figure 15:**
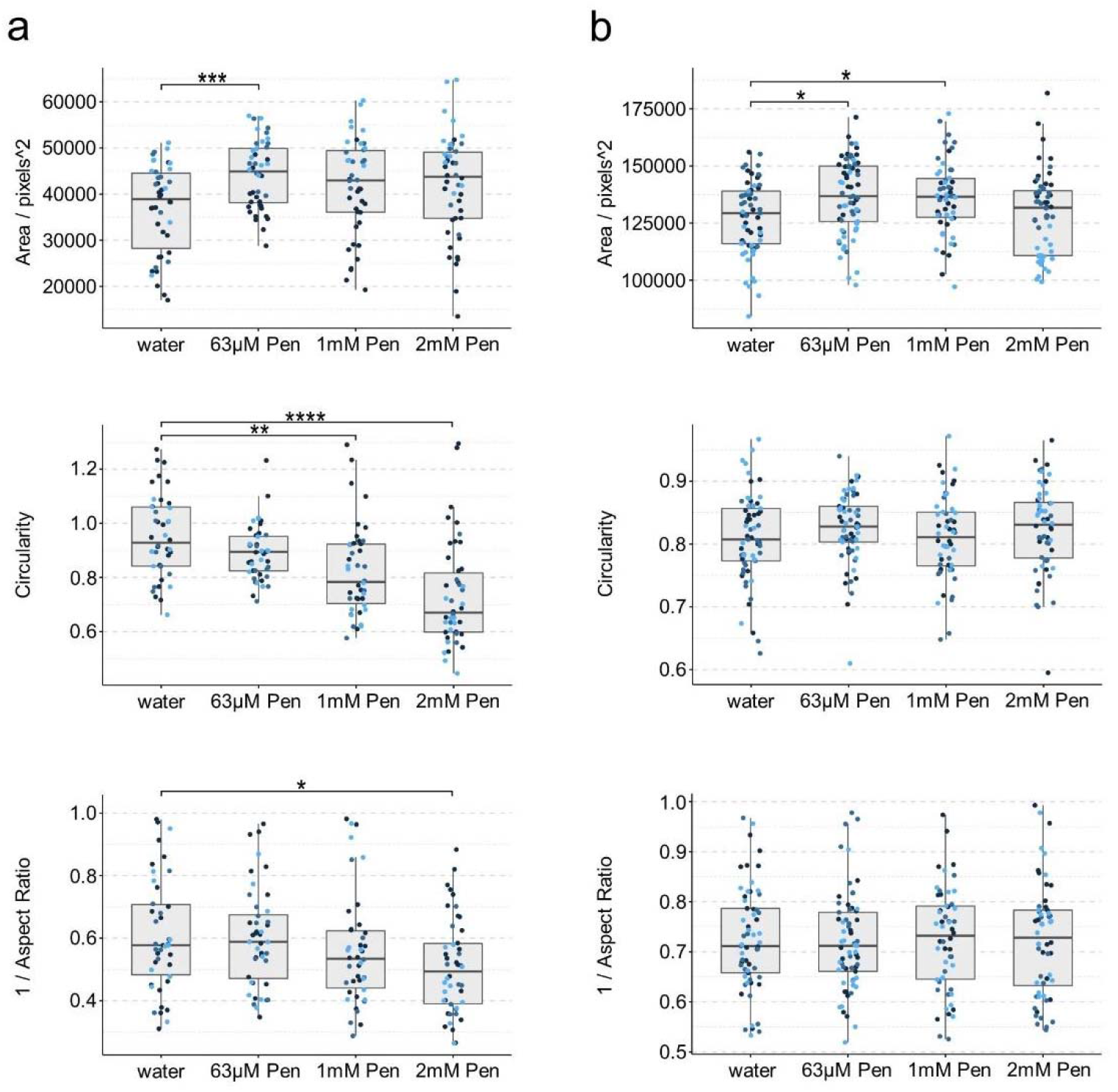
Gastruloid quantification following penicillin G exposure. Quantification of morphology of mouse gastruloids (a), and human gastruloids (b) including Area (top), Circularity (middle) and 1/Aspect Ratio (bottom). Dot colors indicate experimental replicates, and boxplots indicate spread of the data. Significant differences between water (vehicle control) and penicillin G (Pen) treatment conditions are indicated in the plots (Student’s t-test, Bonferroni corrected. Adjusted p-values less than 0.05 are indicated by asterisks (see Materials and Methods for thresholds)).

### 3.8. Gastruloid Patterning Disruption

In summary, retinoic acid and valproic acid appear to be similarly teratogenic in both the mouse and the human gastruloid systems. The human system also identifies ±-thalidomide and bosentan as potentially teratogenic on the basis of altered patterns of reporter gene expression. It would seem that ibuprofen has a more subtle effect in the 63-250 μM range, with both systems pointing towards a possible adverse effect at higher concentrations and altered patterns of gene expression across this range. Finally, our results suggest that the range of phenytoin concentrations that were tested falls below the limit of detectable effect in the human system, inviting its reinclusion in any follow-up study (Tab.2). The T/Bra::GFP reporter line in the mouse suggests that even these low concentrations of phenytoin may disrupt gene expression.

In general, the observed effects are consistent with the results of our previous study on hiPSC-derived EBs [18]. When examining a comparable range of concentrations, we detected the inhibition of cardiomyocyte differentiation with the mEST and a human hiPSC-derived assay measuring quantitative changes in early developmental gene expression (Tab.3, Suppl. Fig.S2). The small panel of reference compounds examined in this study showed variable concentrationdependent effects on gastruloids, with some interesting species-specific differences. Importantly, these differences could be quantified using our image analysis pipeline to identify statistically significant changes in morphology. Together these values can be used to develop a simple readout metric of the assay. Further specification of a defined threshold would enable binary cut-offs to be drawn from the continuous data output, that might provide valuable metrics for a generalized assay design (Fig.16).

**Figure 16:**
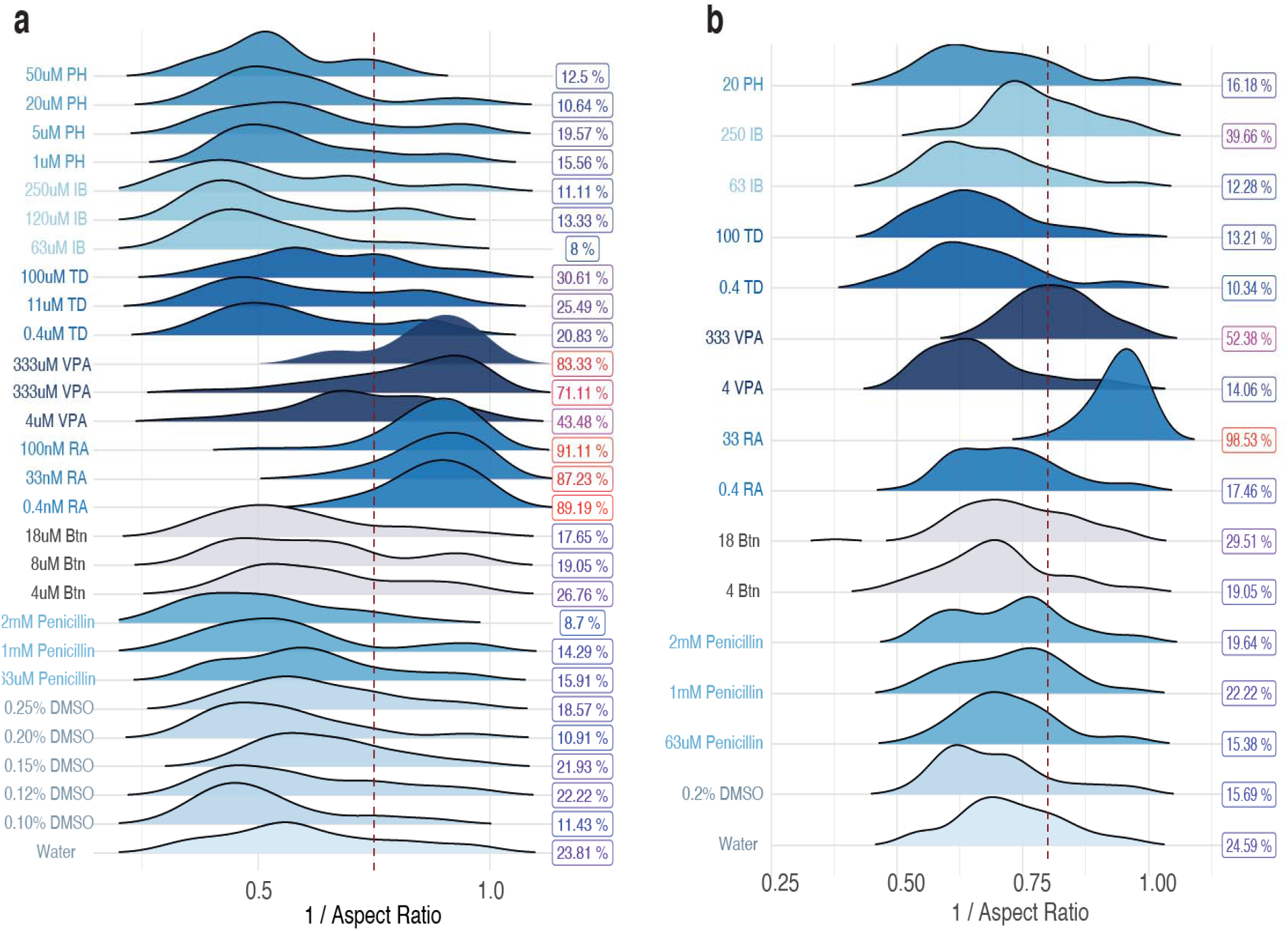
Using quantitative morphological features as a threshold criteria. Density plots of mouse gastruloids (a) and human gastruloids (b) following exposure to each compound. Color indicates treatment compound. Arbitrary 1/Aspect Ratio thresholds of 0.75 for mouse and 0.8 for human gastruloids are drawn (dotted red lines). Percentages (right) of gastruloids above this threshold are shown, and colored by proportion. Btn, Bosentan; RA, Retinoic Acid; VPA, Valproic Acid; TD, Thalidomide; IB, Ibuprofen; PH, Phenytoin.

## 4. Discussion

From this small proof-of-concept study, we conclude that gastruloids might represent a useful tool to assess the effect of compound exposure (in a concentration-dependent manner) on cell differentiation, viability and tissue morphology in a developmental context. This is a particular advantage of using a 3D embryo-like model system with spatiotemporal organization of gene expression, since the gastruloids have intrinsic patterning that mirrors elements of the gastrulating embryo [13, 15, 24]. Although embryoid body systems reflect cellular differentiation into all three germ layers, the gastruloid systems involve the additional asset of spatiotemporal organisation and morphological rearrangement [18, 25, 26]. Here we were able to combine these embryo-like gastruloid models with a toxicological context by assessing morphological perturbations following chemical exposure with multiple phenotypic readouts.

Further experimentation with a wider range of concentrations, a greater number of compounds and different dosing regimens would be required to definitively examine the ability of the gastruloids to provide a comprehensive assay for *in vivo* prediction of teratogenicity or toxicity. An extension to the work in this manner would allow quantitative estimates of assay specificity, sensitivity and accuracy to be identified, which are currently beyond the scope of a study of this size.

However, the results identified here allow for tentative comparisons to be made with existing datasets from animal models or comparative *in vitro* assays such as EB differentiation. When examined in comparison with such datasets, the gastruloids appear to be able to distinguish between known teratogens (such as valproic acid and retinoic acid) and those that are less teratogenic (ibuprofen) or non-teratogenic (penicillin G) [27–33]. They are even able to recapitulate known species-specific sensitivities such as the human-specific effect of thalidomide [3, 4, 34] and can highlight subtle changes in developmental gene expression (bosentan, thalidomide, phenytoin).

It is clear that there is a range of informative outputs from the assay that describe the effect of the compound, including: morphological shape changes (circularity, lack of elongation, size effects - as seen with lower concentrations of retinoic acid in the mouse system), gene expression effects (such as the decrease in *SOX2* expression in human gastruloids following exposure to higher concentrations of valproic acid, which interferes with neurogenesis), and cytotoxicity effects (such as those seen with exposure to high concentrations of retinoic acid or valproic acid in mouse gastruloids) [35]. This suggests that the system can be used to distinguish non-specific effects on cell viability and replication from more specific changes to patterns of gene expression or axial elongation morphogenesis. Together these provide a reference framework of different levels of observable effect that can be used to gauge the response of the system (see Table 2).

Interestingly, we also observed that the human gastruloid model has shown to be more sensitive regarding compound exposure in comparison to the mouse gastruloids. This could be due to species-specific effects upon exposure to certain teratogens. Alternatively, the higher sensitivity of the human gastruloids might also be due to a greater informational content of the data by using triple reporters as readout. Different effects might also be caused by different duration of compound exposures to the mouse gastruloids (120 h) compared to the human gastruloid (72 h). However, this observation is corroborated by results from previous *in vitro* studies, performed with mouse and human embryoid body based systems like the mEST and the hiPSC-derived EB model with extended treatment durations, where human EBs tend to be more sensitive to exposures compared to the murine EBs (Tab.3).

All teratogens were correctly identified with the human gastruloid model whereas the mouse model only predicted adverse morphological effects for retinoic acid and valproic acid. The mouse model predicted adverse effects on gene expression from ±-thalidomide, phenytoin and ibuprofen with less pronounced morphological changes. The negative reference compound penicillin G did not cause any adverse effects in either model. The lowest observed adverse effect concentration levels where morphological changes were obtained with the gastruloid models were consistently comparable to the inhibitory concentrations of existing mouse (ID50) and human *in vitro* models (TC_20_, Tab.3, Suppl. Fig.S2).

In this context, it would be necessary to compare the LOAEL concentrations obtained by both gastruloid models with *in vivo* parameters to set up translational extrapolation models for pharmacokinetics [36, 37]. With the limited range of tested concentrations, we observed three to six-fold lower LOAEL concentrations of the gastruloids compared to *in vivo* Cmax data.

Retinoic acid was an exception, which was detected as positive with more than 2,500-fold lower LOAEL concentrations in the mouse gastruloids compared to rat LOAEL. Cmax and human therapeutic plasma concentrations were also about 3,000-fold higher than the LOAEL concentration of human gastruloids (Tab.3). However, it is necessary to consider that for some compounds, there is a high plasma protein binding (e.g. valproic acid ≥90%, phenytoin 90%). Thus, free fractions of unbound compound (e.g. VPA ~0.3 - 0.5 mM, phenytoin ~3 – 9 μM) reflect human therapeutic concentrations more appropriately, when they are set into correlation with free in vitro concentrations of the human gastruloid model, since the E6 human gastruloid medium is free of serum albumin [38–40].

Given the fact that the gastruloids are grown in 96-well plates and assessed using a plate-based imaging system, there is significant scope for automation and increased throughput in gastruloid generation, imaging and analysis if required. Moreover, implementation of an integrated cytotoxicity assessment to predefine concentration ranges that do not cause impaired cell viability but still induce teratogenic effects, would be beneficial for cytotoxic ranges that are not well described. This would subsequently allow for quantitative assessment additional to size determination or circularity of the gastruloids [41, 42]. The setup of quantitative thresholds for cytotoxicity and morphological dysregulation will be crucial for the establishment of a comprehensive prediction model.

Towards this end, close examination of the proportion of gastruloids under a threshold value of elongation (arbitrary threshold used for demonstration purposes), gives a quick metric output that could serve as a proxy for morphological effects following compound exposure. Such values could be used to design a simple yes/no pass criteria for a given compound exposure and provide proof-of-concept for the assay (Fig. 16). These thresholds can be determined empirically by contrasting known teratogens to known non-teratogens. Establishing this classification framework will enable the unbiased classification of compounds with unknown teratogenic status and it would be a valuable extension of an automated analytical workflow.

## 5. Conclusion

With this study, we have shown that the gastruloid system represents a useful tool for the determination of teratogenic effects during development. Based on a reference panel of seven different compounds we have described morphological and gene expression changes dependent on compound exposure. We were able to differentiate between positive and negative outcomes and even detected species-specific effects. As such, gastruloids represent a novel and potentially powerful assay for teratogenic exposure that utilises the embryo-like spatiotemporal organisation and morphological structure in both mouse and human cell systems. Future efforts could include an extended panel of compounds from different chemical classes with known and unknown teratogenicity, and could establish a clear, quantitative metric for teratogenic and cytotoxic effects as a predictive classifier.

## 6. Ethical Statement

The human gastruloid model used in this study does not show any evidence of cell types associated with anterior neural fates, which would be required to form brain tissue, nor do they form extra-embryonic tissues, which would be required for implantation, or show evidence of multi-organ differentiation, which would be necessary for integrated organ system development. Notably, they lack the morphology of an early human embryo, and therefore do not manifest human organismal form. As such, they are non-intact, non-autonomous, and non-equivalent to *in vivo* human embryos, and do not have human organismal potential. Our research was subject to review and approval from the Human Biology Research Ethics Committee of the University of Cambridge, in compliance with the ISSCR 2016 guidelines.

## 7. Acknowledgments

The human part of this work was supported by grants from Medical Research Council (MRC) (MR/R017190/1 and MR/V005367/1) to A.M.A and N.M, a Leverhulme Trust grant (RPG-2018-356) to AMA, as well as a Gates Cambridge Scholarship awarded to V.M. The mouse part was supported by an ERC Advanced investigator grant (AdG 834580) to AMA supporting PBB. Data derived from human iPSC *in vitro* model based compound assessment has been supported by the BMBF, EFSA, and the DK-EPA (MST-667-00205) and received funding from the European Union’s Horizon 2020 research and innovation program under grant agreements No. 681002 (EU-ToxRisk) and No. 825759 (ENDpoiNTs). The Authors would also like to thank Nicole Schäfer for her valuable contributions in generating all the mEST related data and Paul Barrow for his generous help and useful scientific discussions.

## 10. Supplementary Figures

**Suppl. Figure S1:**
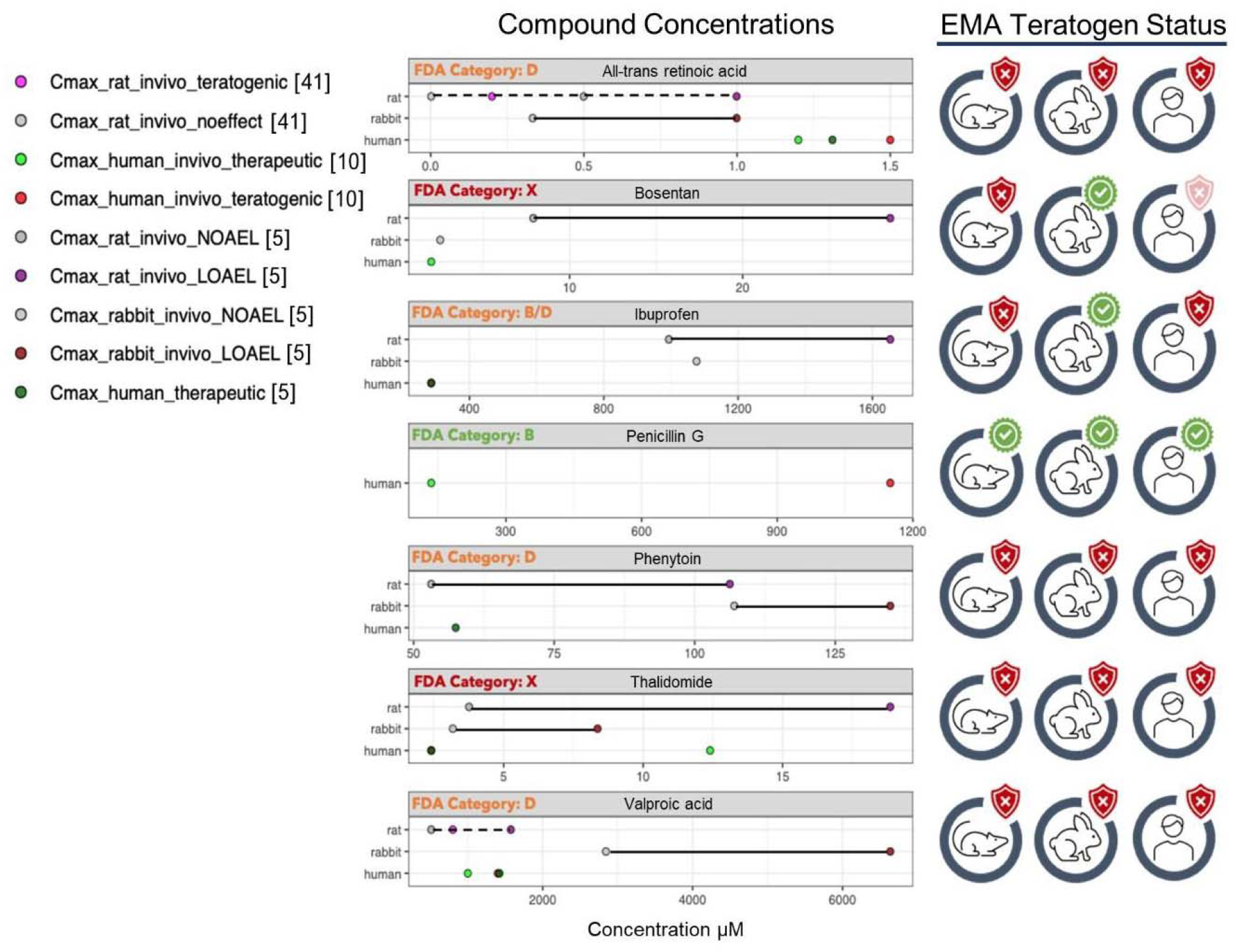
*In vivo* exposure assessment of reference compounds. Using data from the literature [5, 10, 41], the concentrations at which No Observable Adverse Effect (NOAEL) and Lowest Observable Adverse Effect (LOAEL) were identified in different species can be seen in the graphic, alongside their given status as a teratogen as identified by the EMA ICH S5 (R3) guideline [5]. The Cmax concentration, at which therapeutic value is observed in humans, is also plotted, to provide an indication of likely exposure levels *in vivo*.

**Suppl. Figure S2:**
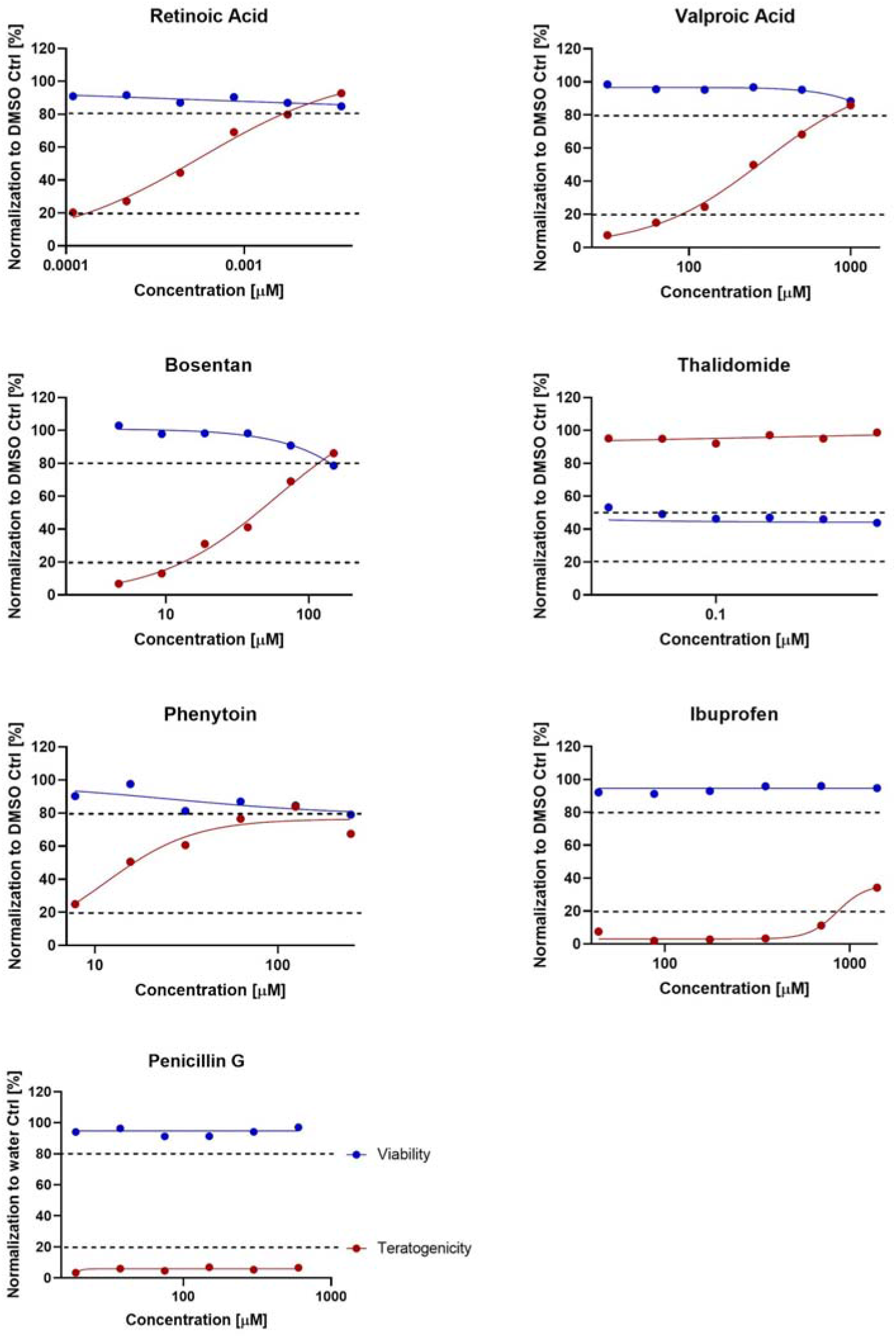
Concentration Response curves for treatments of hiPSC-derived EBs. The graphs represent compound concentration dependent responses of viability (blue) up to a maximum accepted cytotoxicity threshold of 20% viability inhibition and concentration response curves of teratogenicity levels (red) induced by differential gene expression of representative early developmental markers [18]. The effective teratogenicity concentration (TC_20_) is determined by a minimal threshold of 20% differential gene expression compared to DMSO or water (penicillin G) as vehicle control.

**Suppl. Figure S3:**
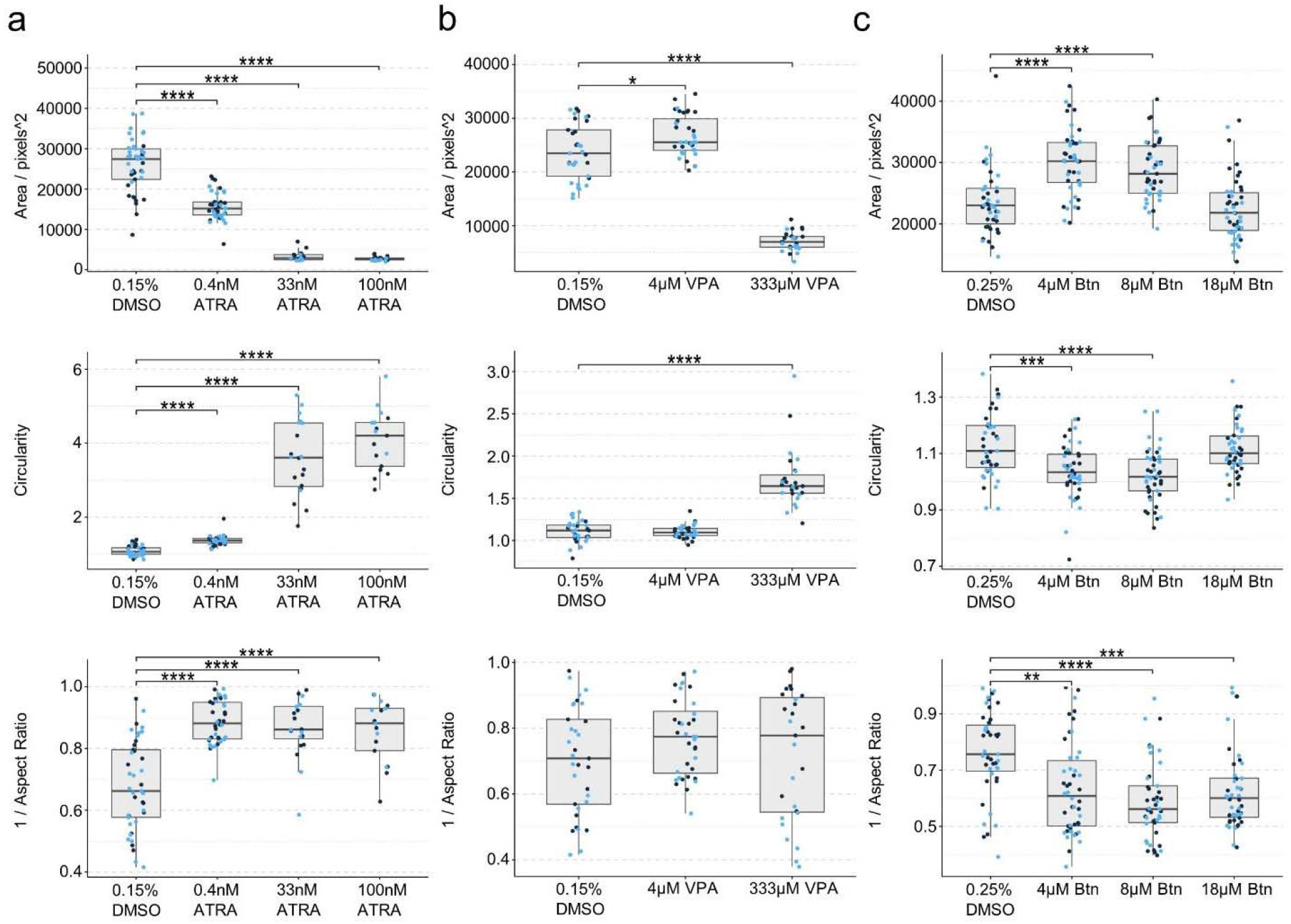
T/Bra::GFP mouse gastruloid quantification following retinoic acid, valproic acid and bosentan exposures. (a-c) Quantification of morphology of T/Bra::GFP mouse gastruloids including Area (top), Circularity (middle) and 1/Aspect Ratio (bottom). Dot colors indicate experimental replicates, and boxplots indicate spread of the data. Significant differences between DMSO (vehicle controls) and *all-trans* retinoic acid (ATRA) (a), valproic acid (VPA) (b), or bosentan (Btn) treatment conditions are indicated in the plots (Student’s t-test, Bonferroni corrected. Adjusted p-values less than 0.05 are indicated by asterisks (see Materials and Methods for thresholds)).

**Suppl. Figure S4:**
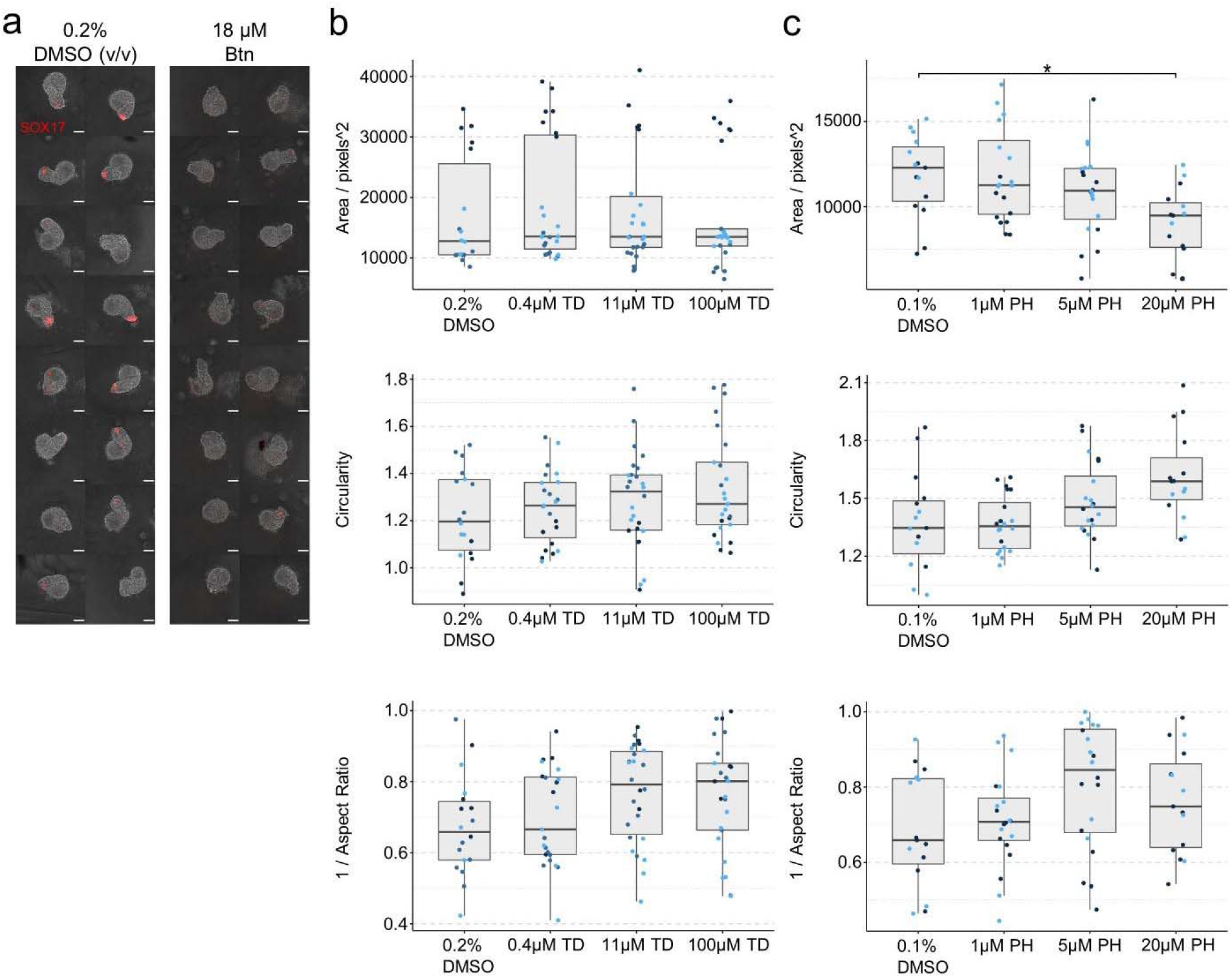
Effect of bosentan treatment on *SOX17* expression in human gastruloids and T/Bra::GFP mouse gastruloid quantification following thalidomide and phenytoin exposures. (a) RUES2-GLR human gastruloids at 72h, following exposure to DMSO (vehicle control) or 18μM bosentan (Btn). Color indicates fluorescent expression of SOX17-tdTomato (red). Scale bars 100μm. (b-c) Quantification of morphology of T/Bra::GFP mouse gastruloids including Area (top), Circularity (middle) and 1/Aspect Ratio (bottom). Dot colors indicate experimental replicates, and boxplots indicate spread of the data. Significant differences between DMSO and thalidomide (TD) (b) or phenytoin (PH) (c) treatment conditions are indicated in the plots (Student’s t-test, Bonferroni corrected. Adjusted p-values less than 0.05 are indicated by asterisks (see Materials and Methods for thresholds)).

**Suppl. Figure S5:**
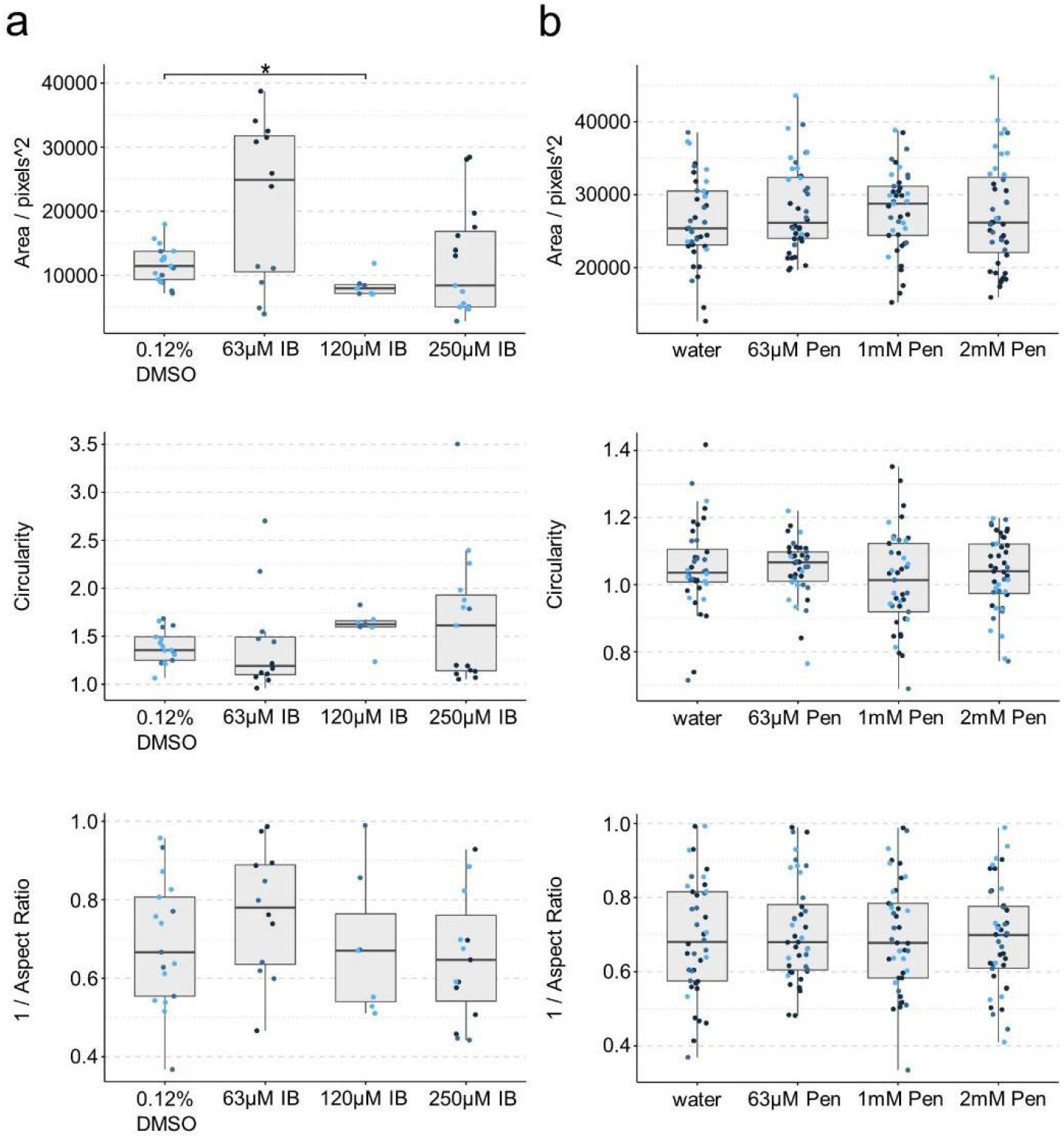
T/Bra::GFP mouse gastruloid quantification following ibuprofen and penicillin G exposures. (a-b) Quantification of morphology of T/Bra::GFP mouse gastruloids including Area (top), Circularity (middle) and 1/Aspect Ratio (bottom). Dot colors indicate experimental replicates, and boxplots indicate spread of the data. Significant differences between DMSO or water (vehicle controls) and ibuprofen (IB) (a) or penicillin G (Pen) (b) treatment conditions are indicated in the plots (Student’s t-test, Bonferroni corrected. Adjusted p-values less than 0.05 are indicated by asterisks (see Materials and Methods for thresholds)).

